# High evidence targets of conserved plant microRNAs and the complexities of their ancient microRNA binding-sites

**DOI:** 10.1101/2022.09.07.507021

**Authors:** Gigi Y. Wong, Anthony A. Millar

**Affiliations:** Division of Plant Science, Research School of Biology, The Australian National University, Canberra ACT 2601, Australia

## Abstract

In plants, high complementarity between microRNAs (miRNAs) and their target genes is a prerequisite for a miRNA-target interaction (MTI). However, evidence suggests there are complexities beyond complementarity that impacts the strength of the MTI. To explore this, the bioinformatic pipeline TRUEE (Targets Ranked Using Experimental Evidence) was applied to strongly conserved miRNAs to identity their high evidence (HE) targets across species. For each miRNA family, HE targets predominantly consisted of homologues from one conserved target gene family (primary family). If an additional HE target family(s) was identified (secondary family), it was likely functionally related to the primary family. Many primary target families contained highly conserved nucleotide sequences flanking their miRNA binding-sites that were enriched in HE homologues across species, suggesting these sequences facilitate miRNA-mediated regulation. A subset of these flanking sequences are predicted to form conserved RNA secondary structures that preferentially base-pair with the miRNA binding-site, implying that these sites are highly structured. Functional testing of the conserved flanking sequences of the miR160 target, *AUXIN RESPONSE FACTOR 10* (*ARF10*), found that mutations within these flanking sequences resulted in attenuated *ARF10* silencing. Our findings support the notion that features beyond complementarity at highly conserved miRNA binding-sites underpin these ancient MTIs.

## INTRODUCTION

The most reported plant microRNAs (miRNAs) in the literature correspond to a set of several dozen miRNAs families that are highly conserved across land plants (Axtell and Meyers, 2018). From nearly two decades of study, it is clear that each of these miRNA families targets a single family of genes via highly conserved miRNA-binding sites (Schwab et al., 2005; Jones-Rhoades, 2012; Tang and Chu, 2017). Underpinning the conservation of these miRNA-target interactions (MTIs) is that many are involved in core biological processes, such as fundamental developmental processes (e.g., miR156, miR160, miR165/166, miR172) (Mallory et al., 2005; Jones-Rhoades, 2012), or environmental responses (e.g., miR395, miR397, miR398) (Sunkar et al., 2006; Abdel-Ghany and Pilon, 2008; Kawashima et al., 2009). As the identity of these conserved target families are predominantly regulatory genes such as transcription factors and F-box proteins, these miRNAs regulate entire gene expression programs (Jones-Rhoades et al., 2006), and reflecting this, perturbation of many of these MTIs result in mutant phenotypes with pleiotropic defects (Todesco et al., 2010).

It has long been known that plant MTIs require a high degree of complementarity (Rhoades et al., 2002; Schwab et al., 2005; Liu et al., 2014). Based on this, bioinformatic tools that predict miRNA targets are based on miRNA-target complementarity. Although this approach has successfully identified most conserved targets (Rhoades et al., 2002; Jones-Rhoades and Bartel, 2004), the degree of complementarity does not strictly correlate with whether a target is strongly or weakly regulated by any given miRNA. Therefore, complementarity alone is insufficient in identifying which predicted target genes are subjected to physiologically relevant miRNA-regulation, implying factors other than complementarity are involved. Given the evolutionary age of these conserved MTIs, it is feasible that additional complexities in MTI have arisen.

Currently, the evidence supporting the existence of such complexities is not clear. Several studies have raised the possibility that miRNA-binding sites are present in highly accessible regions of the target transcripts. Firstly, a bioinformatic analysis across multiple species found AU rich synonymous codons are enriched in nucleotides (nts) that flank miRNA-binding sites (Gu et al., 2012). This correlated with a greater miRNA-binding site accessibility via the selection for reduced RNA secondary structures. Supporting this, a study on the RNA secondary structure of the Arabidopsis transcriptome found the 21 nt miRNA-binding site to be less structured compared to the 50 nt sequences immediately flanking upstream and downstream of these miRNA-binding sites (Li et al., 2012). However, as this was an *in vitro* study, conclusions drawn must be taken in the context that it was conducted in the absence of cellular influences. Indeed, opposing this study, a recent *in vivo* study found miRNA-binding sites to be highly structured, with their unfolding being the limiting factor of cleavage efficiency directed by a miRNA-induced silencing complex (miRISC) (Yang et al., 2020). Here, only the two nucleotides immediately downstream of the miRNA-binding site were required to be single stranded for efficient cleavage (Yang et al., 2020).

Supporting the notion of a highly structured miRNA-binding site, was the discovery of highly conserved RNA secondary structures associated with the miR159-binding site of two *GAMYB* genes in Arabidopsis (*MYB33* and *MYB65*). These RNA structures were functionally demonstrated to making these genes highly sensitive to miR159-mediated silencing, independent of AU content or predicted miRNA-binding site accessibility, highlighting our lack of understanding (Zheng et al., 2017). Likewise, a predicted RNA stem-loop lies within a conserved region flanking the miR399-binding site of *INDUCED BY PHOSPHATE STARVATION1* (*IPS1*) (Franco-Zorrilla et al., 2007; Wong et al., 2018). *IPS1* has been used as a miRNA target *MIMIC* (Todesco et al., 2010), but with variable efficacy (Reichel et al., 2015). It was shown that strengthening the stem of this RNA secondary structure could strongly enhance the efficacy of the *MIM* when targeting miR165/166, make RNA secondary structure a likely determinant of *MIM* efficacy (Wong et al., 2018). This all argues that RNA secondary structure associated with miRNA-binding sites can strongly impact MTIs.

Given the long evolutionary history of these ancient MTIs, we have hypothesized that additional regulatory mechanisms beyond miRNA-target complementarity have arisen that contribute to miRNA specificity (Li et al., 2014). It is apparent that these conserved MTIs are fixed across species, with multiple members of a specific miRNA family regulating multiple members of a specific target family (Li et al., 2014; Axtell and Meyers, 2018). Given the dominance of these highly conserved target families, we previously hypothesized that they could be considered as the primary target(s) of these conserved miRNA families, and that as an active miRISC is all that is needed to execute silencing, the acquisition of any additional targets would be constrained by the requirements of the primary miRNA-target relationship. Finally, driving the functional specificity of miRNAs, we hypothesized that complexities beyond complementarity may be limiting the promiscuity of functional miRNA-targeting (Li et al., 2014).

In this paper, these hypotheses are explored. Firstly, we apply our recently published miRNA target pipeline, Targets Ranked Using Experimental Evidence (TRUEE) targets (Wong and Millar, 2022), to the set of highly conserved miRNAs across diverse plant species. TRUEE assesses the strength and frequency of degradome signals across multiple libraries for each predicted target to identify High Evidence (HE) and Low Evidence (LE) targets. These identified HE targets of conserved miRNAs were then examined to identify any potential conserved features beyond complementarity that may be contributing to the functional specificity of these miRNAs.

## RESULTS

### HE targets primarily consist of a single gene family for most conserved miRNA families

TRUEE was applied to 20 highly conserved miRNAs and a tasiARF across diverse plant species to identify HE and Low Evidence (LE) targets. As expected, LE targets outnumbered HE targets for most miRNAs across species (Figure 1). The exception was miR169, where LE targets consisted of half the total predicted targets. For miRNAs in which the expectation score was increased from 3.0 to 5.0, the number of LE target increased by almost an order of magnitude.

**Figure 1.**
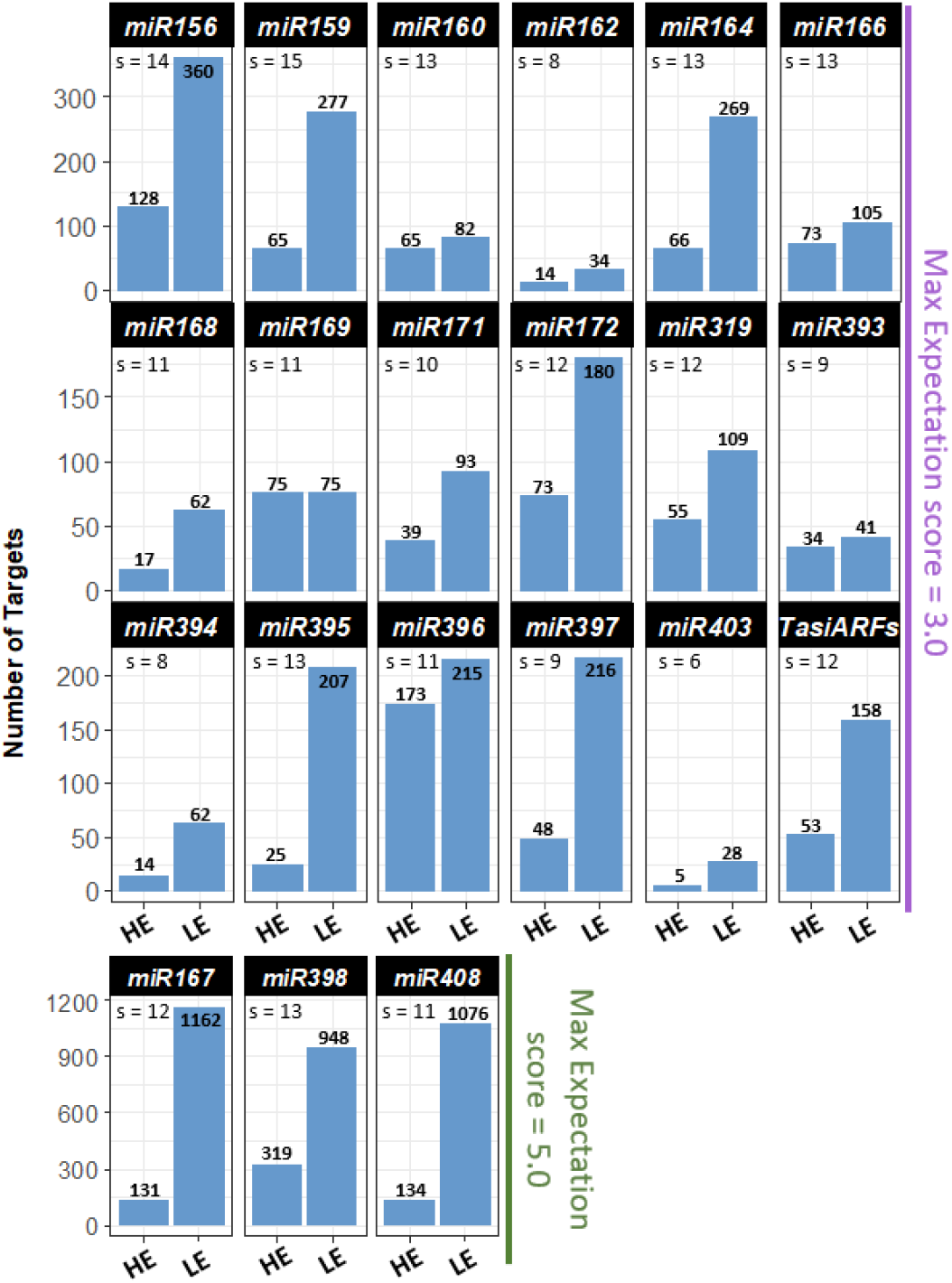
HE and LE targets for conserved miRNAs from diverse species. The number of HE and LE target genes across species for 20 conserved miRNAs and the tasiARF. Numbers on the top of bars indicate the number of genes found across all species. s = number of species analysed per miRNA.

To determine the number of gene families targeted by each conserved miRNA family, the gene family of HE and LE targets was identified using their associated PANTHER ID (Protein ANalysis THrough Evolutionary Relationships) (Mi et al., 2013). Results show that for each miRNA and tasiARF, HE targets across species were predominantly composed of homologues of the same gene family (Figure 2). These families were the same as those most often reported in literature to be targets of their corresponding miRNA, and hence, considered here as the primary target family (Jones-Rhoades and Bartel, 2004; Jones-Rhoades, 2012; Sunkar et al., 2012; Chauhan et al., 2017). For some miRNAs, HE targets are almost exclusively made up of homologues of this primary target family; miR160 [94% - *AUXIN RESPONSE FACTOR* (*ARF*)], miR166 [99% - *CLASS III HOMEODOMAIN LEUCINE ZIPPER* (*HD ZIPIII*)], miR170/miR171 [97% - *HAIRY MERISTEM* (*HAM*)], miR172 [95% - *APETELA2-LIKE* (*AP2*)] and tasiARF (98% - *ARF*). Therefore, these miRNAs-target relationships appear fixed across species.

**Figure 2.**
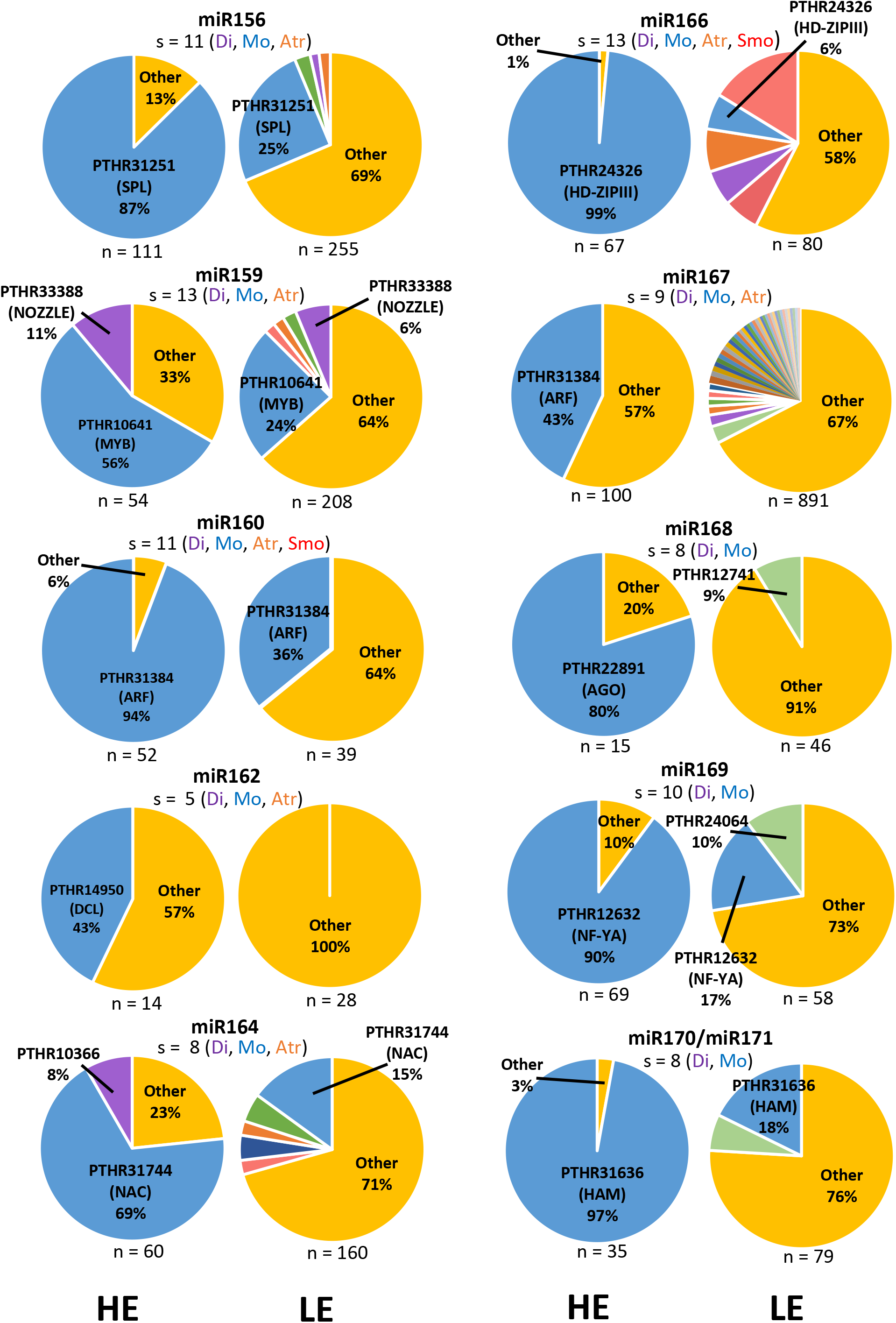
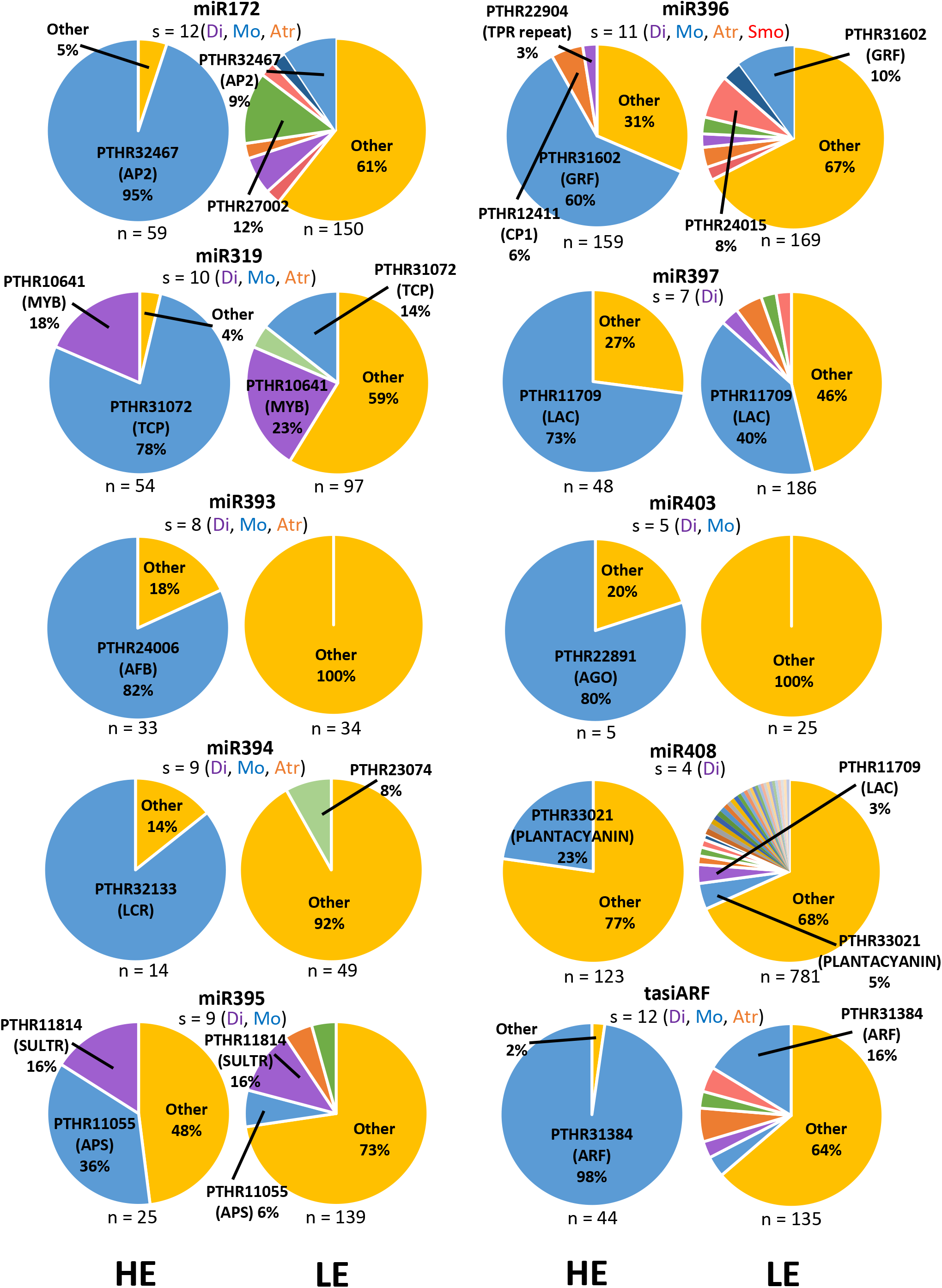
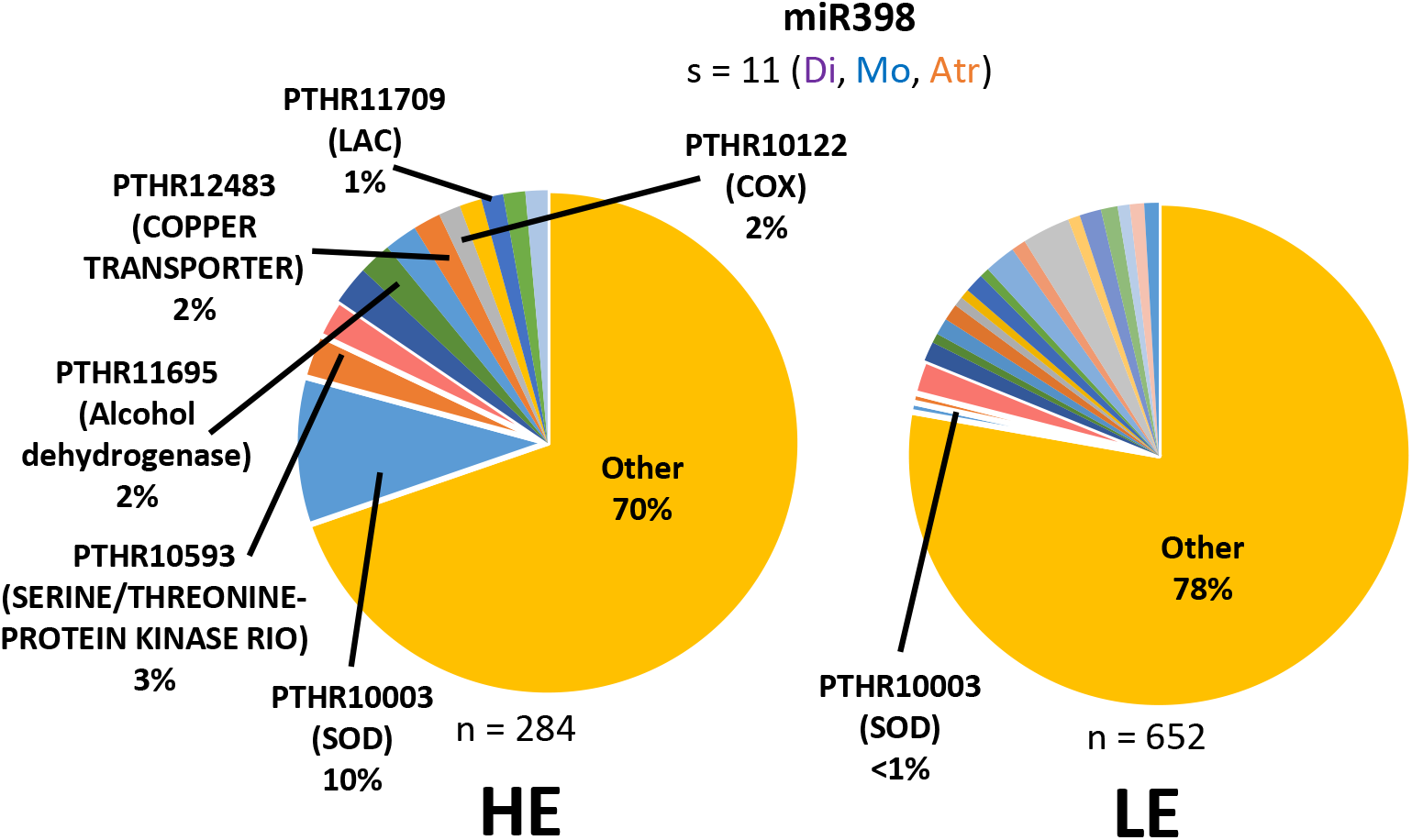
Distribution of the gene families of HE and LE targets per sRNA. HE and LE targets were categorised into gene families by their associated PANTHER ID. PANTHER IDs with 3 or less members were grouped into the ‘other’ category. n = the number of HE or LE targets used in the analysis. s = the number of species used in the analysis. Di = dicots; Mo = monocots; Atr = *A. trichopoda;* Smo = *S. moellendorffii*. The PANTHER ID and gene family name is indicated. Targets with no associated PANTHER ID are not included in the analysis, hence the total number of targets is less than in Figure 1.

In contrast, LE targets predominantly consisted of genes from diverse PANTHER IDs (Figure 2). This is indicated in that they largely consisted of gene families which are grouped in the ‘other’ category (PANTHER IDs associated with three or less members). In all but one miRNA family (miR397), the ‘other’ category made up over half of the total LE targets. Furthermore, in all cases, the percentage of primary target families in LE targets were smaller than in the HE targets. Strikingly, for some miRNA families, no or very few primary target family members are found to be LE targets; this includes miR162, miR166, miR167, miR168, miR393, miR394, miR398 and miR403 (Figure 2). In fact, no primary target family members were identified as LE targets for miR394; only one for miR162; and two for miR167 and miR393. This highlights the prevalence to which these primary target families are subjected to miRNA-mediated regulation.

### The few secondary HE target families identified are less conserved and have less members

For the HE targets, six miRNA families (miR159, miR164, miR319, miR395, miR396 and miR398) regulate additional conserved target families (defined as having four or more conserved HE targets across multiple plant species) (Figure 2). These additional HE target families, henceforth called secondary target families, had fewer HE targets compared to the primary target families and were found in a narrower range of species (Table 1). For instance, whereas HE targets from all primary target families were found beyond dicotyledonous species, HE targets from the secondary target families were restricted to dicotyledonous species with the exception of miR319:*MYB*. However, miR319 is closely related to miR159 and can also target *MYB* genes, although in Arabidopsis, it was shown that targeting of *MYB* by miR319 is limited, with miR159 being the major regulator (Palatnik et al., 2007). This also appears conserved across species, as miR319-mediated regulation of *MYB* genes is much weaker than the corresponding miR159-mediated regulation as indicated by their respective target plots (T-plots) (Figure S1). Therefore, the general trend remains; a conserved miRNA family predominantly regulates one primary target family that is conserved across species, and although acquisition of secondary target families occurs, targeting of these families is less conserved and fewer homologues are regulated.

**Table 1.** Distribution of HE target family members across species. The number of HE targets from the primary target family (in bold) and secondary target families found in each species. The species analysed are; *Arabidopsis thaliana* (At), *Citrus sinensis* (Cs), *Glycine max* (Gm), *Malus domestica* (Md), *Medicago truncatula* (Mt), *Prunus persica* (Pp), *Solanum lycopersicum* (Sl), *Vitis vinifera* (Vv), *Brachypodium distachyon* (Bd), *Hordeum Vulgare* (Hv), *Oryza sativa* (Os), *Zea mays* (Zm), *Amborella trichopoda* (Atr), and *Selaginella moellendorffii* (Sm).

|  |  | At | Cs | Gm | Md | Mt | Pp | Sl | Vv | Bd | Hv | Os | Zm | Atr |
| --- | --- | --- | --- | --- | --- | --- | --- | --- | --- | --- | --- | --- | --- | --- |
| **miR159** | **PTHR10641 (MYB)** | **2** | **2** | **6** | **1** | **2** | **1** | **3** | **2** | **1** | **4** | **2** | **2** | **2** |
|  | PTHR33388 (NOZZLE) | 1 |  | 3 |  |  | 1 | 1 |  |  |  |  |  |  |
| **miR164** | **PTHR31744 (NAC)** | **5** | **1** | **10** | **4** | **3** | **2** | **3** | **3** | **4** |  | **3** | **1** | **2** |
|  | PTHR10366 (NAD<br>DEPENDENT<br>EPIMERASE/DEHYDRATASE) |  |  | 1 |  |  | 1 | 3 |  |  |  |  |  |  |
| **miR319** | **PTHR31072 (TCP)** | **5** |  | **10** | **8** | **2** | **2** | **4** | **2** |  |  | **4** | **4** | **1** |
|  | PTHR10641 (MYB) |  |  | 6 |  |  |  |  | 2 |  |  | 1 | 1 |  |
| **miR395** | **PTHR11055 (APS)** | **2** | **1** |  | **2** |  | **1** | **2** | **1** | **1** |  |  |  |  |
|  | PTHR11814 (SULTR) |  |  | 3 |  |  |  |  | 1 |  |  |  |  |  |
| **miR396** | **PTHR31602 (GRF)** | **6** | **6** | **24** |  | **9** | **6** | **10** |  | **9** |  | **11** | **9** | **6** |
|  | PTHR12411 (CP1) | 2 | 1 | 3 |  | 1 |  | 2 |  |  |  |  |  |  |
|  | PTHR22904 (TPR repeat) |  |  | 3 |  | 1 |  |  |  |  |  |  |  |  |
| **miR398** | **PTHR10003 (SOD)** | **3** | **1** | **6** | **4** | **1** | **2** | **1** | **2** | **1** |  | **3** |  | **3** |
|  | PTHR10122 (COX) | 1 |  | 2 |  | 2 |  |  | 1 |  |  |  |  |  |
|  | PTHR11709 (LACCASE) |  |  |  |  | 1 | 2 | 1 |  |  |  |  |  |  |
|  | PTHR33021 (PLANTACYANIN) | 1 |  |  |  | 2 |  |  | 1 |  |  |  |  |  |
|  | PTHR12483 (COPPER<br>TRANSPORTER; COPT) |  |  | 2 | 3 |  | 1 |  |  |  |  |  |  |  |
|  | PTHR10593 (SERINE/THREONINE-PROTEIN KINASE RIO;<br>STPKR) |  |  | 6 |  |  |  |  | 2 |  |  |  |  |  |
|  | PTHR11695 (ALCOHOL<br>DEHYDROGENASE; ADH) |  |  | 5 |  | 1 | 1 |  |  |  |  |  |  |  |

### Target families of the same miRNA are commonly functionally related

Supporting the hypothesis that a secondary target would need to be compatible with the primary miRNA-target relationship (Li et al., 2014), three of six proposed secondary miRNA families were from functionally related processes to the primary target family. For instance, functional studies in Arabidopsis have shown that for the miR159 target families, *MYB* and *NOZZLE*, both are involved in anther development (Schiefthaler et al., 1999; Millar and Gubler, 2005); the miR395 targets, *ATP-SULFURYLASE* (*APS*) and *SULFATE TRANSPORTER2;1* (*SULTR2;1*), are both involved in sulfur metabolism and transport (Liang et al., 2010); and the miR398 targets, *SUPEROXIDE DISMUTASE* (*SOD*) and *CYTOCHROME C OXIDASE* (*COX*), are both involved in response to oxidative stress (Sunkar et al., 2006; Yamasaki et al., 2007) (Table 1). Furthermore, for miR398, *LACCASE* (*LAC*) and *PLANTACYANIN*, which are copper proteins like *SOD* were also identified as secondary target families (Abdel-Ghany and Pilon, 2008). An additional copper transporter gene family (*COPT*; PTHR12483) was also identified as a secondary target family of miR398 in dicots outside of Arabidopsis (Table 1) (Naya et al., 2014). Together, this suggests that in the instances in which a secondary target family is acquired, they are likely from functionally related processes.

### Complementarity is not an absolute determinant of HE targets across miRNAs

Previous miRNA target prediction programs have relied heavily on the ranking of targets by miRNA-target complementarity (Bonnet et al., 2010; Sun et al., 2011; Dai et al., 2018). However, it is unclear how strict the correlation is, as there are targets with 3-5 mismatches that are strongly miRNA-regulated, whilst there are genes with 0-2 mismatches which are poorly regulated (Brousse et al., 2014; Liu et al., 2014; Zheng et al., 2017).

Therefore, to investigate the extent that miRNA-target complementarity can be used as an indicator of HE targets, the complementarity of all HE and LE targets was calculated via their Expectation Score, a psRNATarget parameter that reflects complementarity (Dai et al., 2018). For most miRNAs, the average Expectation Scores were generally lower for the HE targets compared to the LE targets (Figure 3). For some miRNA targets, [e.g., miR160, miR166, miR171, miR394, miR403 and tasiARF], HE targets had a much lower average Expectation Score compared to LE targets, suggesting complementarity is a strong determinant. However, the average Expectation Scores were not statistically different for all miRNAs [eg. miR162, miR319, miR395, and miR408] which suggests that it is not a reliable indicator for all miRNA families.

**Figure 3.**
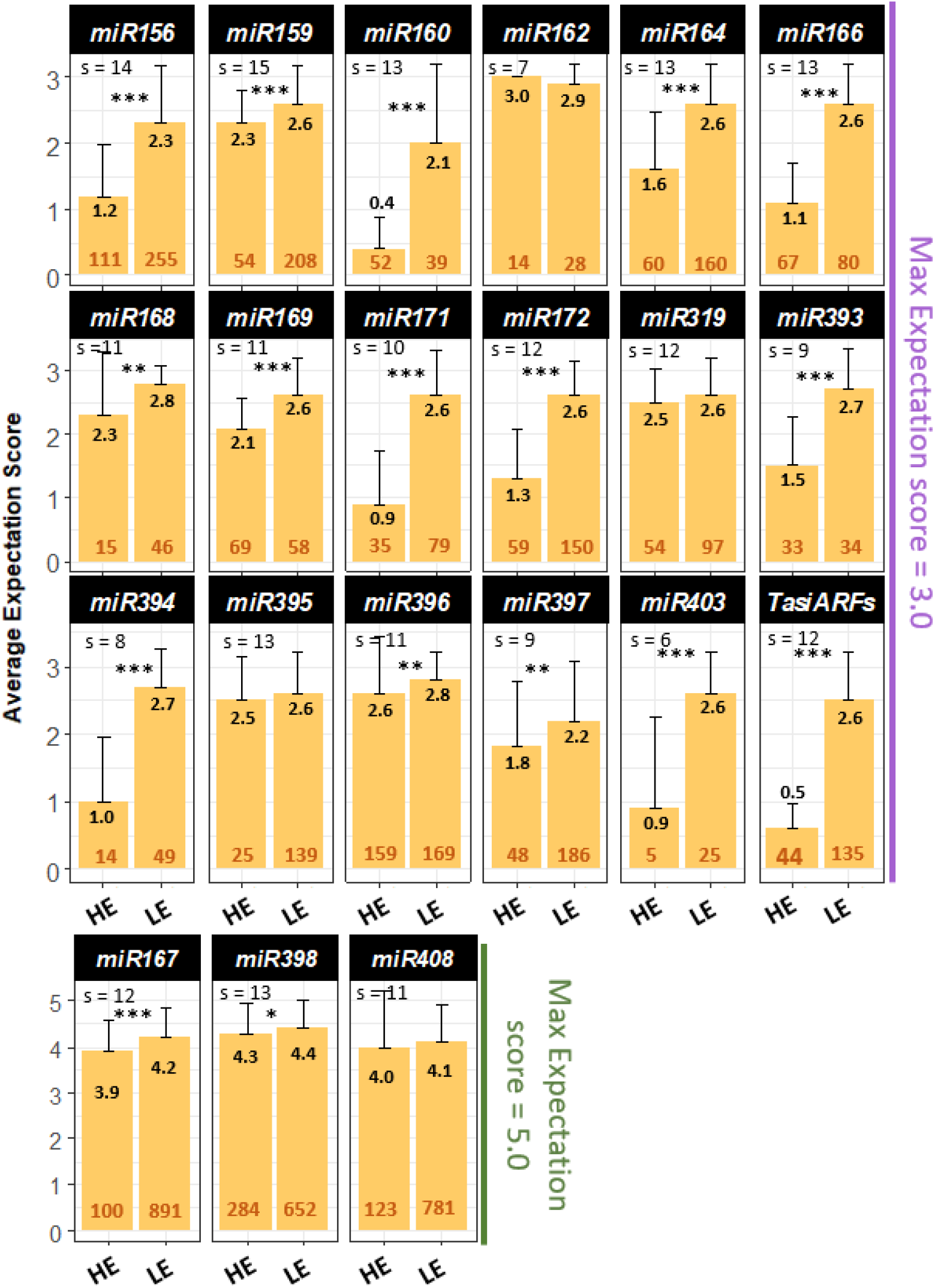
The average expectation score for HE and LE targets across species for each sRNA. The expectation score for target prediction with psRNATarget was increased to 5.0 (green) for miR167, miR398 and miR408 as the known targets for Arabidopsis exceed an expectation score of 3.0. Expectation score for targets for all other miRNAs do not exceed 3.0 (purple). s = the number of species used for each analysis. Orange numbers = the number of genes analysed. Asterisks = statistical difference between the Expectation scores of HE and LE targets where * indicates P ≤ 0.05, ** indicates P ≤ 0.01, *** indicates P ≤ 0.001 and no asterisks indicates no significant difference.

These results suggest that miRNA-target complementarity requirements vary greatly between different miRNA families. As such, ranking the confidence of a gene as a miRNA target based on Expectation Score cannot be generally applied across miRNA-target pairs. This implies factors additional to miRNA binding-site complementarity are involved in the MTI.

### Many miRNA target families have conserved sequences flanking their miRNA binding-sites

Previously, we showed that highly conserved nucleotides flanking the miR159-binding site of GAMYB homologues that corresponded to the stems of ancient RNA secondary structures (Zheng et al., 2017; Millar et al., 2019). Moreover, it was functionally demonstrated that these RNA secondary structures facilitated miR159-mediated silencing of *MYB33* in Arabidopsis (Zheng et al., 2017). Therefore, we hypothesized that similar scenarios may have arisen in other highly conserved target families over evolutionary time. To investigate this, multiple sequence alignments (MSAs) were performed on the primary target families, followed by phyloP analysis to identify whether nucleotide conservation extends beyond their miRNA-binding site (Hubisz et al., 2011).

#### miR160:*ARF10*

*ARF10* homologues were identified in multiple dicot and monocot species and twenty were used to construct a MSA (Figure 4A). The miR160 binding site was invariant in all but four homologues. Conservation extended six nucleotides beyond the miR160 binding site at both the 5‵ and 3‵ ends. Both these flanking sequences contained four nts that are complementary to the miR160-binding site, hence potentially forming a conserved RNA secondary structure within the miR160-binding site (Figure 4B).

**Figure 4.**
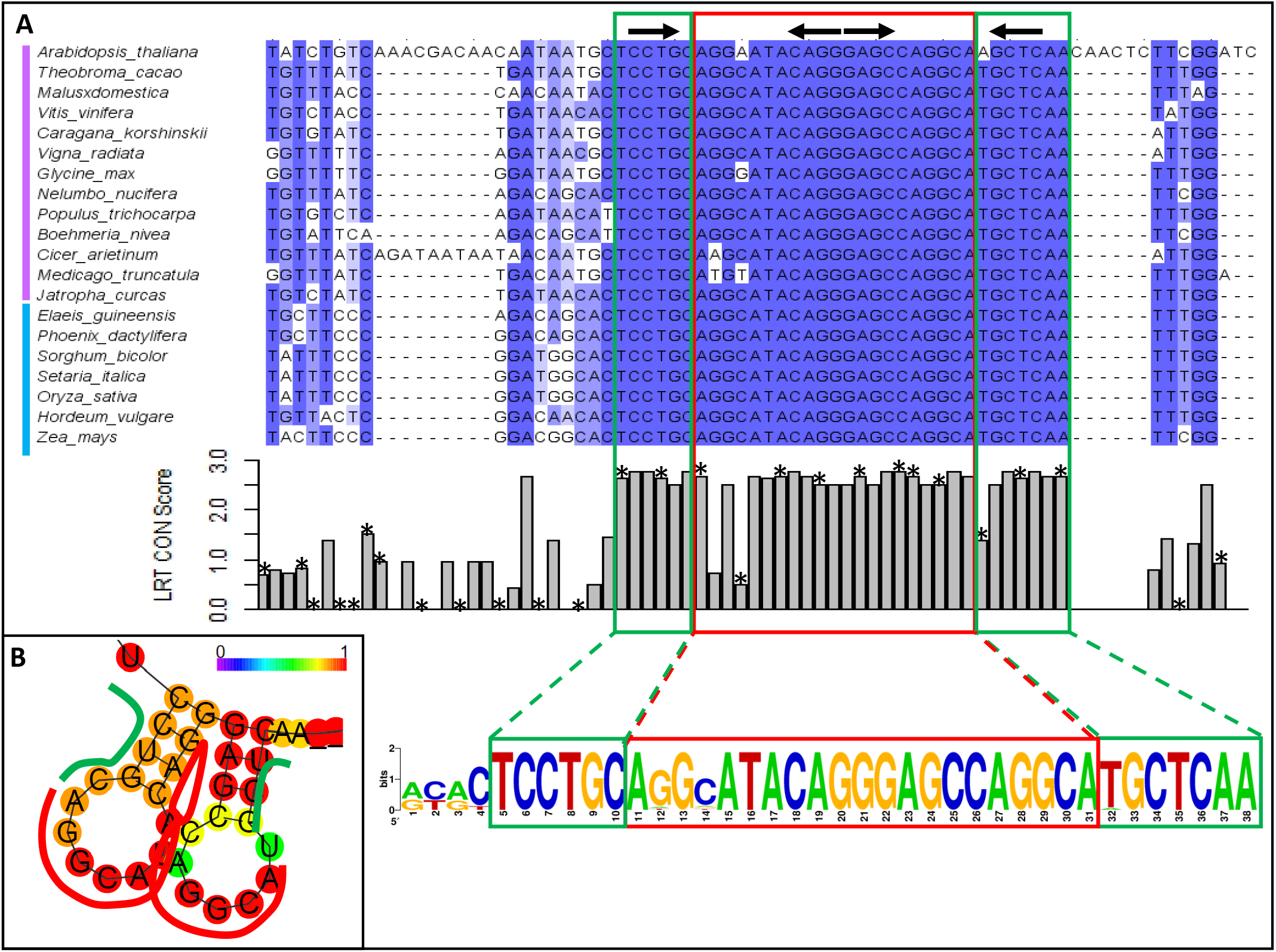
Conserved nucleotides flanking the miR160-binding site of ARF10 homologues. **(A)** A multiple sequence alignment (MSA) of twenty ARF10 homologues from various dicot (pink bar) and monocot (light-blue bar) species. The miR160 binding-site is indicated by a red box, and the conserved flanking sequences in green boxes. The phyloP score at each nucleotide position around the binding site is indicated, where * = a degenerate nucleotide site in reference to the Arabidopsis sequence. Black arrows above the MSA indicate base pairing from the predicted RNA secondary structure in **(B)** The RNA secondary structure and probability of base pairing predicted from the consensus sequence from the MSA.

#### miR160:*ARF17*

*ARF17* homologues were identified from dicots, monocots, the basal angiosperm, *A. trichopoda*, lycophytes, and to the oldest extant lineage of land plants, bryophytes. Aside from the 5‵-nucleotide position, the miR160 binding site was invariant in all but three homologues and twenty were used to construct a MSA (Figure 5A). Conservation extended to seven flanking nucleotides directly upstream of the miR160 binding site and was near-identical in homologues across these diverse lineages, suggesting it corresponds to an ancient motif. Interestingly, all seven conserved flanking nucleotides are complementary to the miR160 binding site and are predicted to form a conserved secondary structure at a high base-pair probability (Figure 5B).

**Figure 5.**
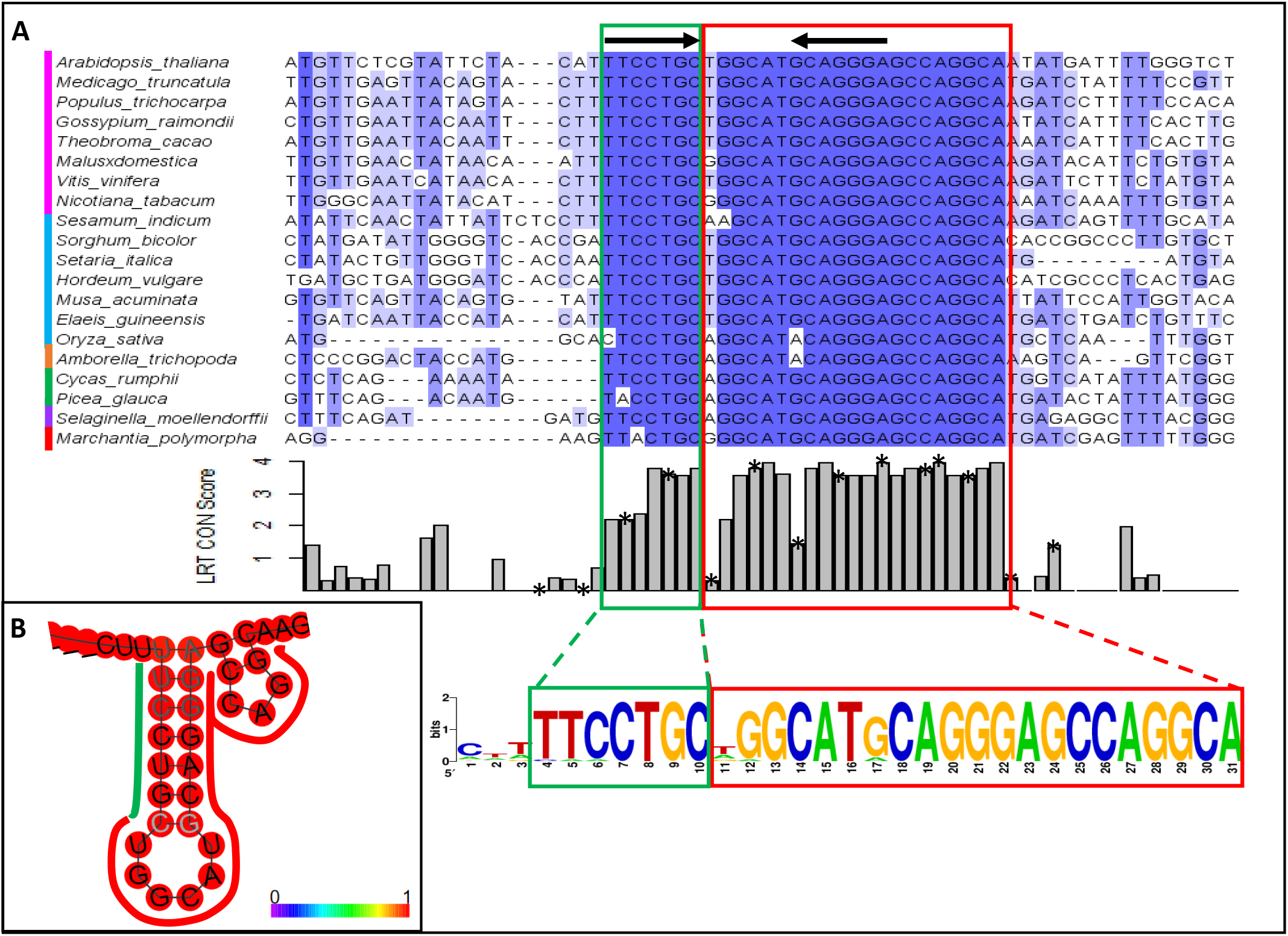
Conserved nucleotides flanking the miR160-binding site of ARF17 homologues. **(A)** A MSA constructed from twenty ARF17 homologues from various lineages of land plants (dicots, monocots, A. trichopoda, gymnosperms, lycophytes, and bryophytes), with the sequence logo of the miR160 binding-site (red box) and conserved flanking sequences (green box). The phyloP score at each nucleotide position around the binding site is indicated, where * = degenerate nucleotide site in reference to the Arabidopsis sequence. Black arrows above the MSA indicate base pairing from the predicted RNA secondary structure in **(B)** The RNA secondary structure and the probability of base pairing predicted from the consensus sequence from the MSA.

#### miR171:*HAM1*

*HAM1* homologues [also known as *SCARECROW-like* (*SCL*)] were identified from multiple dicot species and twenty were used to construct a MSA (Figure 6). *HAM1* has two overlapping binding sites (miR171b, c, and miR170/miR171a) that are offset by three nucleotides (Bari et al., 2013). No nucleotide variation was found in either binding site across the homologues analysed. The sequences directly flanking these binding sites were highly conserved, five nucleotides upstream and nine nucleotides downstream (Figure 6A). These conserved flanking sequences were predicted to form the stems of RNA secondary structures. Like *ARF10* and *ARF17*, the conserved sequence directly upstream of the miRNA binding sites was predicted to base pair with a GC rich sequence in the binding site to form a strong stem (Figure 6B).

**Figure 6.**
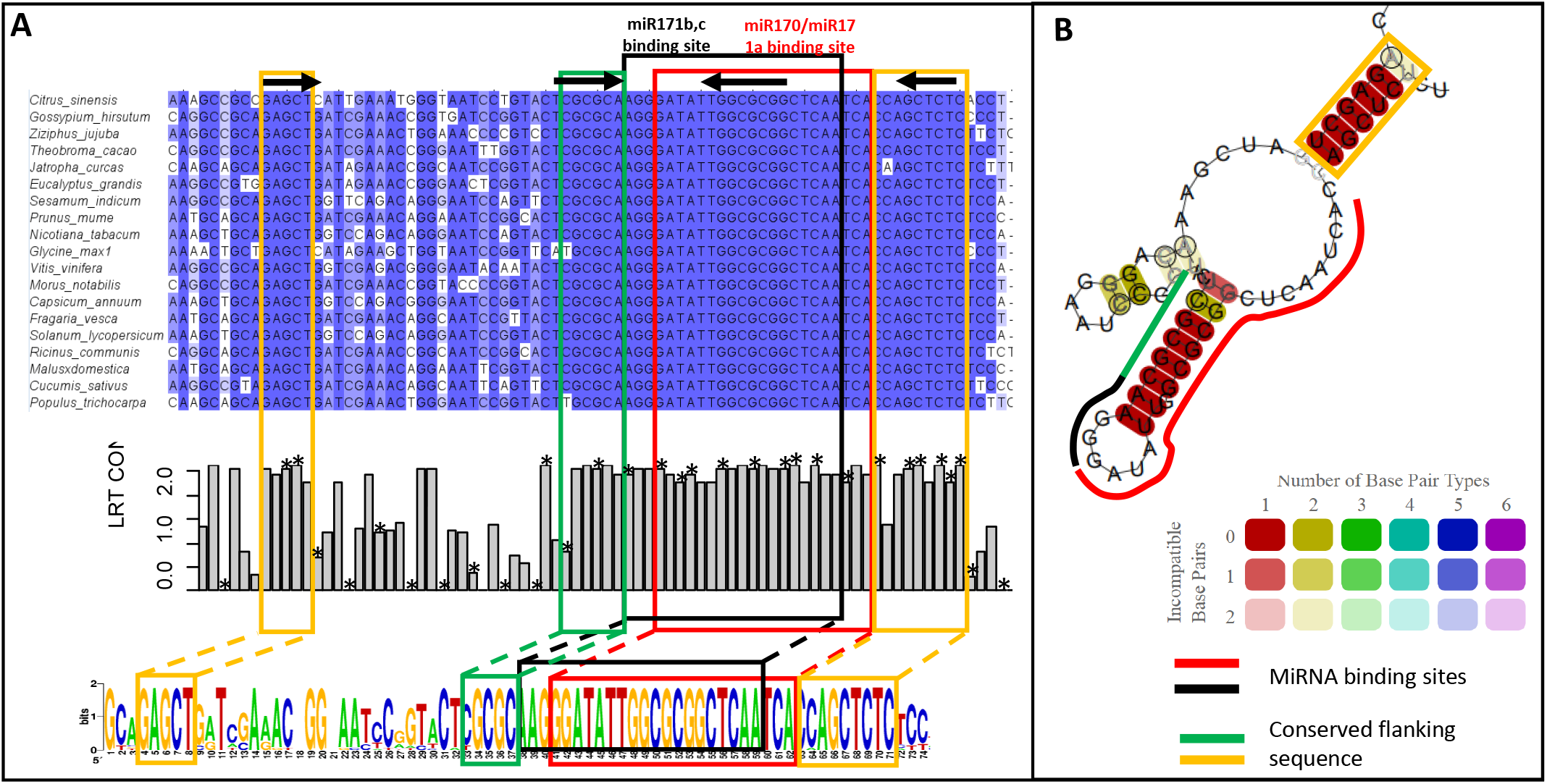
Conserved nucleotides flanking the miR171-binding site of HAM1 homologues. **(A)** A MSA constructed from twenty HAM homologues from various dicot species. HAM1 homologues have two miRNA-binding sites which overlap by three nucleotides. The miR171b,c binding site is indicated by a black box, and the miR170/miR171a in red. The conserved flanking sequences are indicated by green and yellow boxes. The phyloP score at each nucleotide position around the binding site. * = a degenerate nucleotide site in reference to the Arabidopsis sequence. Black arrows above the MSA indicate base pairing from the predicted secondary structure in **(B)** The RNA secondary structure and the probability of base pairing predicted from the consensus sequence from the MSA.

#### miR319:*TCP2*

Twenty *TCP2* homologues were identified in multiple dicot and monocot species and used to construct an MSA (Figure S2). Nine nucleotides were found to be conserved upstream of the miR319 binding site, five of which corresponded to wobble positions, suggesting conservation at the RNA level. This conserved sequence was not predicted to form an RNA secondary structure.

#### miR319:*TCP4*

Similarly, for *TCP4*, twenty homologues from multiple dicot and monocot species were used to construct an MSA (Figure S3). In this case, two conserved sequences were found with one conserved sequence consisting of five nucleotides directly flanking the 5‵ end of the binding site. Another sequence was downstream of the binding site and was four nucleotides long. Like *TCP2*, no RNA secondary structure was predicted to form from these sequences.

#### miR390:*TAS3*

For the non-coding RNA, *TRANS-ACTING SHORT INTERFERING RNA 3* (*TAS3*), a MSA of twenty *TAS3* homologues from dicot species found conservation to extend upstream of the miR390-binding site (Figure S4). These consisted of two conserved sequences which were five and nine nucleotides long (from distally to proximally of the binding site, respectively) and were separated by a three-nucleotide gap. These conserved sequences were not predicted to form an RNA secondary structure. However, these sequences have been reported as corresponding to siR1769 which is derived from a phased product (Allen et al., 2005).

#### miR396:*GRF3*

Twenty *GROWTH-REGULATING FACTOR 3* (*GRF3*) homologues from multiple dicots, monocots species and *A. trichopoda* were used to construct a MSA (Figure S5). Three conserved sequences were identified upstream of the miR396 binding site. Conserved sequences were not predicted to form an RNA secondary structure.

#### tasiARF:*ARF2*

Twenty *ARF2* homologues from multiple dicots and monocots were used to construct a MSA (Figure 7A; Figure S6). Conserved sequences consisting of seven nucleotides each were identified upstream and downstream of the tasiARF binding site (Figure 7A). All conserved nucleotides were complementary to the binding site and were predicted to base-pair at high probability to form two stem-loops (Figure 7A). Nucleotide variations were found in one position in the conserved sequences which was still compatible with base-pairing to the binding site (Figure S6).

**Figure 7.**
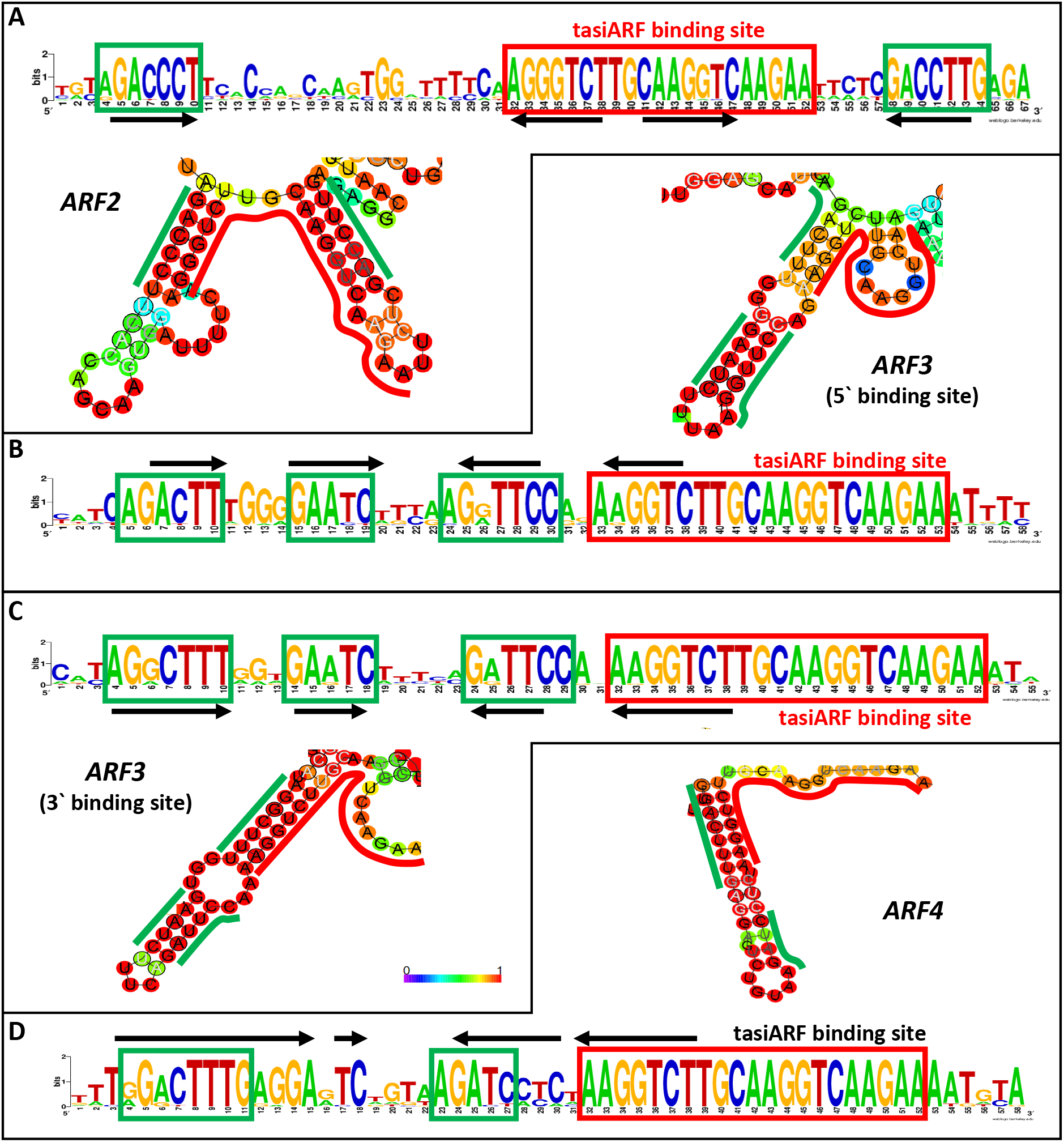
Conserved nucleotides flanking the tasiRNA-binding site of ARF homologues. The sequence logo of the tasiARF binding-site (red box) and conserved flanking sequences (green boxes from various MSAs constructed from twenty ARF homologues. The RNA secondary structure and the probability of base pairing predicted from the consensus sequences for each MSA is shown, with black arrows above the sequence logos indicating base pairing in the predicted RNA secondary structures. **(A)** *ARF2*; **(B)** 5‵ binding-site in *ARF3*; **(C)** 3‵ binding-site in *ARF3*; **(D)** 5‵ binding-site in *ARF4*.

#### tasiARF:*ARF3*

*ARF3* contains two tasiARF-binding sites [see Fahlgren et al. (2006) for schematic]. For the 5‵ binding site a MSA was constructed from *ARF3* homologues from diverse lineages ranging from dicots, monocots, *A. trichopoda* and gymnosperms. Three conserved sequences were found upstream of the binding site across these lineages (Figure 7B; Figure S7) and had sequence complementarity that was predicted to form a stem-loop that incorporated the 5‵ end of the tasiARF binding-site (Figure 7B). Nucleotide variations which were still compatible with base pairing were found at two positions in the conserved sequence in multiple homologues. In one of these positions one variant (G) appeared to be more common in the dicots and monocots, and the another (A) more common in homologues from more ancient lineages (*A. trichopoda* and gymnosperms) (Figure S7).

For the 3‵ binding-site, a MSA was constructed from *ARF3* homologues from the same species as above. Similarly, three conserved sequences were found directly upstream of the binding-site across all lineages (Figure 7C; Figure S8) and had sequence complementarity that was predicted to form a stem-loop that incorporated the 5‵-end of the tasiARF binding-site (Figure 7C). Nucleotide variations which were still compatible with base-pairing were found at two positions in the conserved sequence in multiple homologues. These conserved flanking sequences were highly similar between both binding-sites in *ARF3*, only differing at four nucleotide positions.

#### tasiARF:*ARF4*

Like *ARF3*, *ARF4* also contains two tasiARF binding-sites. For the 5‵ binding-site, a MSA constructed from *ARF4* homologues of twenty dicot and gymnosperm species showed two conserved sequences upstream of the binding site (Figure 7D; Figure S9). The 5‵ conserved sequence was complementary to the 3‵ conserved sequence and the tasiARF binding-site with which it was predicted to form a stem-loop (Figure 7D). Nucleotide variations which were still compatible with base-pairing were found at four positions in the conserved sequence in multiple homologues (Figure S9).

For the 3‵ binding site, a MSA was constructed using twenty homologues from diverse dicot species (Figure S10). These species differed from the ones used in *5‵* binding-site and were chosen as they had the highest conservation while retaining diversity in the species. Two conserved sequences upstream of the binding site was identified which had complementarity to the binding site with which it was predicted to form a stem-loop (Figure S10).

Comparison of the consensus flanking sequences upstream of all the tasiARF binding-sites of *ARF2*, *ARF3* and *ARF4*, found a high degree of sequence identity (Figure S11). Furthermore, any nucleotide variations between these flanking sequences still retained the potential to base-pair with the tasiARF binding-site, being either single nucleotide variations (e.g., an A or G, to base-pair with a U) or compensatory substitutions. This pattern of nucleotide conservation argues selection for an RNA stem is occurring. Conversely, the poorest conservation was found at the nucleotide positions that were predicted to form the loop region. This pattern of conservation is consistent with selection for these distinct RNA secondary structures that encompass the tasiARF binding-site.

### Conserved sequences flanking the miRNA binding-sites are enriched in HE targets

Having identified conserved sequences flanking the miRNA binding-site in multiple target families, it was investigated how these sequences were distributed among HE and LE target homologues using the sequence and spacing criteria (as detailed in Figure S12-S14).

For all target families, a higher percentage of HE targets possessed the conserved sequences compared to LE targets indicating that these sequences are enriched in targets subjected to strong miRNA-mediated regulation (Figure 8). This was most strikingly for miR159, where the conserved flanking sequences were exclusively found in *GAMYB* homologues that corresponded to HE targets (23 of the 30 *GAMYB* HE homologues) (Figure 8). This occurred across diverse species, including the ancient basal angiosperm, *Amborella trichopoda*. This strongly supports the idea that these sequences, which correspond to an ancient RNA secondary structural element, have been central in the miR159-mediated regulation of *GAMYB* targets across species (Table 2).

**Figure 8.**
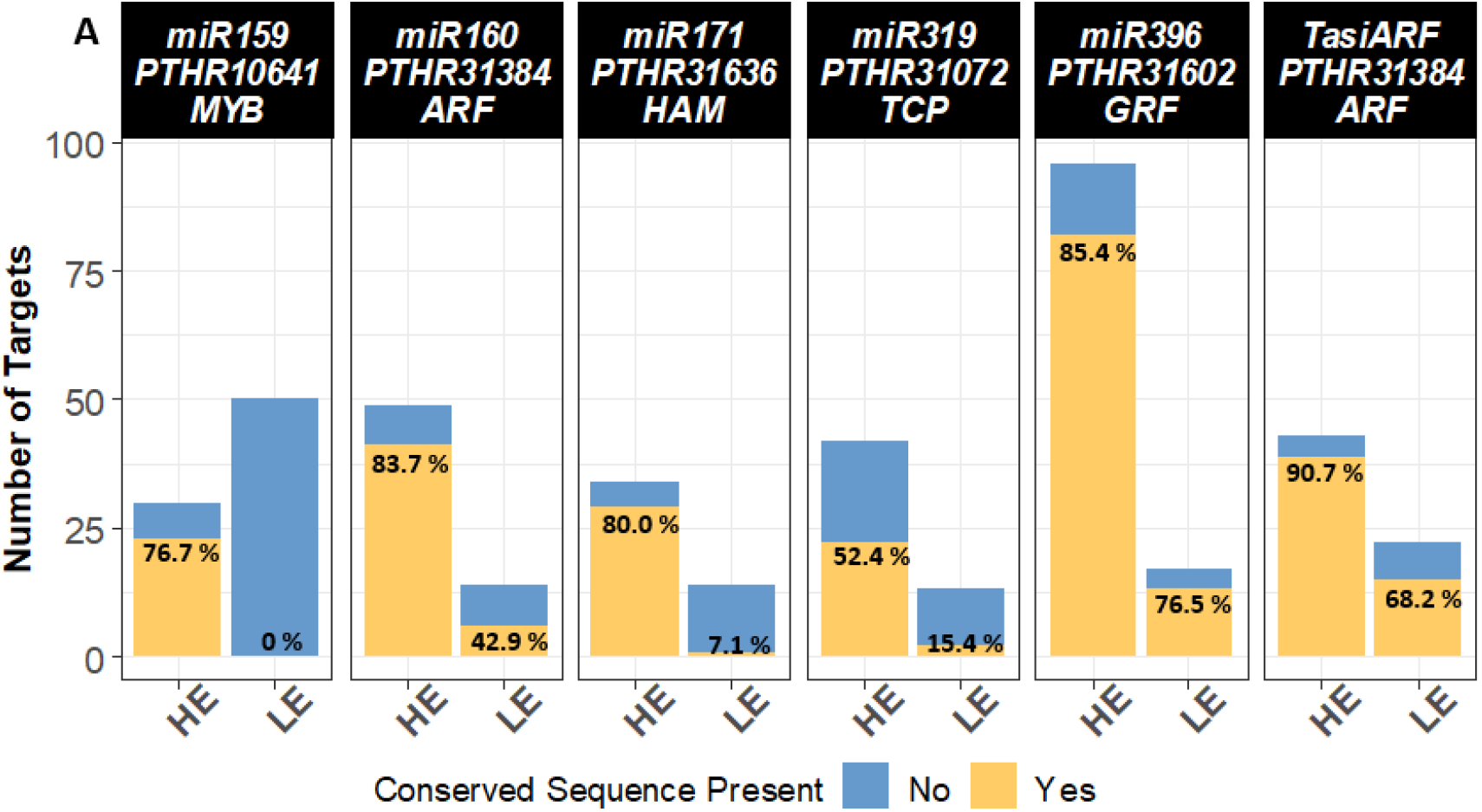
Presence of conserved sequences flanking the binding site in HE and LE targets in conserved MTIs. Genes with conserved sequences are indicated in yellow and genes without conserved sequences are indicated in blue. The number of genes with the conserved sequence out of total genes is indicated as a percentage in the yellow bars.

**Table 2.** The number of HE targets possessing the conserved sequences flanking the miRNA binding-site across species. The species analysed are; *Arabidopsis thaliana* (*At*), *Citrus sinensis* (*Cs*), *Glycine max* (*Gm*), *Malus domestica* (*Md*), *Medicago truncatula* (*Mt*), *Prunus persica* (*Pp*), *Solanum lycopersicum* (*Sl*), *Vitis vinifera* (*Vv*), *Brachypodium distachyon* (*Bd*), *Hordeum Vulgare* (*Hv*), *Oryza sativa* (*Os*), *Zea mays* (*Zm*), *Amborella trichopoda* (*Atr*), *and Selaginella moellendorffii* (*Sm*).

|  | At | Cs | Gm | Md | Mt | Pp | Sl | Vv | Bd | Hv | Os | Zm | Atr | Sm |
| --- | --- | --- | --- | --- | --- | --- | --- | --- | --- | --- | --- | --- | --- | --- |
| **miR159 - MYB** | **2** | **1** | **6** | **1** | **1** | **1** | **2** | **2** | **1** | **3** | **1** | **1** | **1** |  |
| **miR160 - ARF** | **2** |  | **10** |  | **5** | **4** | **3** | **4** | **3** |  | **3** | **4** | **1** | **2** |
| **miR171 - HAM** | **3** | **2** | **6** |  | **2** | **2** | **3** | **3** | **1** |  | **3** | **4** |  |  |
| **miR319 - TCP** | **3** |  | **6** | **5** | **2** | **1** | **2** | **2** |  |  |  |  | **1** |  |
| **tasiARF - ARF** | **3** | **3** | **11** | **3** | **4** | **3** | **2** | **3** | **3** |  | **4** | **3** |  |  |

For all other target families, enrichment of the conserved flanking sequences in HE targets was not as strong, but nevertheless occurred, most notably *HAM* (miR171) and *TCP* (miR319) homologues (Figure 8). In all cases these homologues were from species spanning beyond dicots; the miR159:*MYB* and miR319:*TCP* family module conservation was found to extend to *A. trichopoda*; the miR160:*ARF* family module to the distantly related lycophyte, *S. moellendorrffii* (Table 2). This supports a functional importance of these sequences in miRNA-mediated regulation that is deeply conserved.

### Conserved flanking nucleotides of the miRNA binding-site of *ARF10* impacts miR160-mediated regulation

Next, we functionally tested whether conserved flanking sequences to the miR160 binding-site impact miRNA-mediated regulation using functional genetic approaches *in planta*. *ARF10* was chosen due to its highly conserved flanking sequences which are predicted to form an RNA secondary structure with high confidence. Additionally, plants where miR160-mediated regulation of *ARF10* has been perturbed are well characterized and have an easily distinguishable phenotype (Liu et al., 2007).

Firstly, an *ARF10* transgene with flanking mutations was made (*ARF10-FM*) via the introduction of synonymous mutations into both the 5‵ and 3‵ conserved flanking sequences which also alters the predicted RNA secondary structure (Figure 9A). *ARF10-FM* was compared to a wild-type *ARF10* transgene (*ARF10-WT*) and an *ARF10* transgene with miR160 binding-site mutations rendering it resistant to miR160-mediated regulation (*rmARF10*) (Figure 9A). All *ARF10* transgenes were individually fused to a CaMV 2×35s promoter of pMDC32 and transformed into Arabidopsis. Primary transformants for each variant was then scored for a mutant leaf curl phenotype which has been previously reported in transgenic plants overexpressing a miR160-resistant *ARF10* transgene (Liu et al., 2007). Phenotypic defects were categorized by severity into; ‘No leaf curl’, where plants displayed no leaf curl and were indistinguishable from wild type plants; ‘Weak’, where plants displayed one curled leaf with the abaxial side visible from an aerial view; and ‘Strong’, where plants displayed two or more curled leaves with the abaxial side visible from an aerial view (Figure 9B).

**Figure 9.**
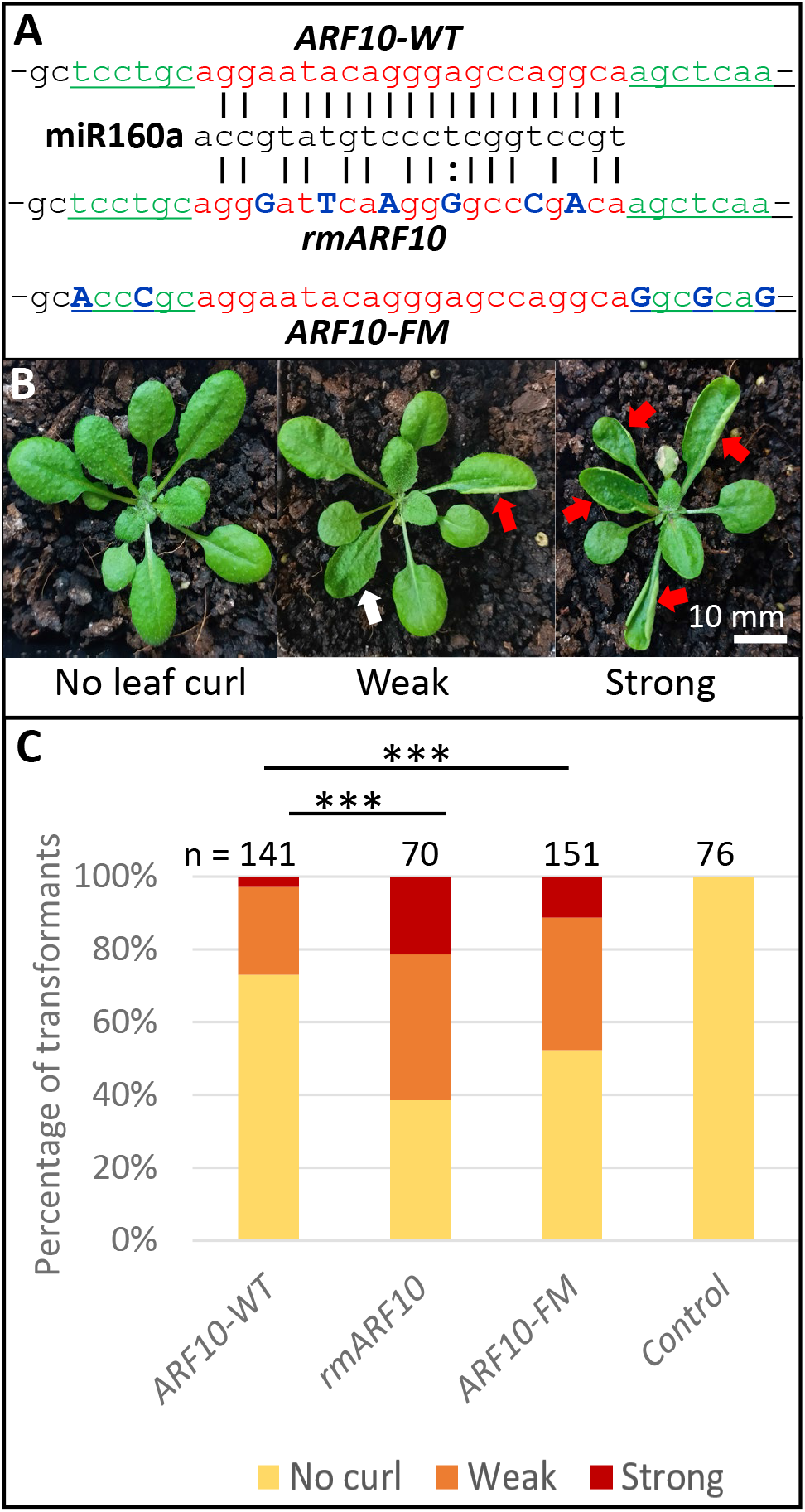
Functional analysis of the conserved flanking nucleotides in *ARF10*. **(A)** Mutations within the *ARF10* gene to result in the *rmARF10* (miR160 binding-site mutations) and *ARF10-FM* (flanking mutations). The miR160 binding-site is indicated in red, and the conserved flanking sequences in green. In *rmARF10* and *ARF10-FM*, mutated nucleotides are blue capital letters. All constructs encode identical amino acid sequences. **(B)** Phenotypes of four-week-old Arabidopsis plants showing different degrees of the mutant rosette phenotype. The red arrows indicate leaves curled so that the abaxial side is showing when viewed from the top. White arrows indicate leaves with apparent leaf curl but without the abaxial side is showing. **(C)** Percentage of *ARF10* primary transformants showing a ‘No leaf curl’, ‘Weak’ and ‘Strong’ phenotype. ‘n’ indicates the number of plants analysed. A vector only transgenic control construct was also included. Both the *rmARF10* (p = 3.63e-04) and *ARF10-FM* (p = 1.85e-07) were significantly different from wild-type.

Results found that most *ARF10-WT* plants displayed a ‘No leaf curl’ phenotype (73%), whereas this was just over half for *ARF10-FM* plants (52%), and even less for *rmARF10* plants (61%). *ARF10-WT* plants also demonstrated less severe phenotypes with only four plants (3%) categorized as having a ‘Strong’ mutant phenotype compared to *ARF10-FM* (11%) and *rmARF10* (21%). Therefore, the conserved sequences flanking the binding site appear to influence miR160-mediated regulation of *ARF10*.

## DISCUSSION

In this study, TRUEE found that each highly conserved miRNA predominantly targeted only a single gene family across species (primary target family), and that targeting a secondary target family was rare. Sequence alignments of homologues of these primary target families found that many contained conserved sequences that flanked their miRNA binding-sites, and that these sequences were predicted to base-pair with the miRNA binding-sites forming conserved RNA secondary structures. Possible functional significance of these features was supported by the fact that they were found to be enriched in gene homologues that were ranked as HE targets by TRUEE, and that mutation of the conserved flanking sequences of the miR160 binding-site in *ARF10* attenuated it regulation by miR160. All these observations argue that conserved MTI have developed complexities that confer a functional specificity that is not apparent from complementarity alone.

### TRUEE demonstrates that the primary conserved MTIs predominate across species

Despite the myriad of predicted MTI, only a small number of these have been experimentally validated. Here, TRUEE demonstrated the extent to which these conserved MTIs predominate across species where for some miRNAs, the identified HE targets were almost exclusively one gene family (eg. miR160:*ARF*; miR166:*HD-ZIPIII*; miR172:*AP2*) (Figure 2). Analysis also failed to identify a miRNA family that switched primary target families; currently the only example in the literature is miR827 (Lin et al., 2018). This contrasts to that of animals, where it is not uncommon for a single miRNA family to target a large number of distinct target families (Buscaglia and Li, 2011; Pfeffer et al., 2015; Chen et al., 2017; Song et al., 2017). Potentially underlying these differences are the complementarity requirement of MTIs in plants, and the strength of the silencing outcomes, both of which seem much higher in plants than in animals.

### Multiple target families of a conserved miRNA are likely to be functionally related

However, TRUEE did find secondary target families for some conserved miRNA families. In many of these instances, the primary and secondary target families were functionally related (Figure 2; Table 1). This was the case for miR159, both *GAMYB* and *NOZZLE* families being involved in anther development (Schiefthaler et al., 1999; Millar and Gubler, 2005); miR395, both *APS* and *SULTR* families being involved in sulfur homeostasis (Liang et al., 2010); and for miR398, which targets multiple families related to copper (Cu), including the primary family *SOD* and secondary target families, *COX*, *LAC* and *PLANTACYANIN* (encode Cu containing proteins; Abdel-Ghany and Pilon, 2008; Sunkar et al., 2006; Yamasaki et al., 2007), and *COPT*, a Cu transporter that is in the same pathway as *SOD* (Pilon, 2017). Other examples in the literature include miR399 that targets the phosphate-regulated genes, *PHOSPHATE2* (*PHO2*), and *IPS1* (Franco-Zorrilla et al., 2007); and miR167 that targets the auxin related genes, *ARF* and *IAA-ALANINE RESISTANT 3* (*IAR3*) genes (Chorostecki et al., 2012).

These observations all support our hypothesis on the constraints of functional MTI in plants (Li et al., 2014). Of the many MTI predicted, only a select subset appear preferentially regulated, and these are predominantly from homologues of a single target family. If targeting of a second conserved target family occurs, it must have a MTI that is compatible to regulatory conditions defined by the primary MTI, and so is likely to be functionally related in some aspect. This occurs as the expression of the miRNA will be dictated by the desired regulatory outcome of the primary target family, and so the regulation of any additional targets must be achieved in that context (Li et al., 2014).

### Complementarity varies greatly between conserved miRNA-target family pairs

It is clear that complementarity requirements varied greatly between each conserved miRNA-target family pair, with the average Expectation Score of HE targets varying from 0.4 for miR160, to 4.3 for miR398 (Figure 3). This implies complementarity cannot be used as a clear indicator of an HE target across miRNA families.

Consistent with this is that Liu et al. (2014) found that binding sites engineered with perfect complementarity to the miRNAs are not the most strongly silenced. Similarly, artificial miRNAs (amiRNAs) engineered with similarly high complementarity to their intended targets also varied in regulation (Deveson et al., 2013; Li et al., 2013). Some miRNA targets with suboptimal complementarities have been experimentally validated, whilst there are other predicted targets with a higher degree of complementarity for which little or no evidence has been found (Debernardi et al., 2012; Brousse et al., 2014). Therefore, this implies that additional factors are involved in the miRNA-mediated regulation of a target.

### A role for RNA secondary structure in facilitating miRNA-mediated regulation?

In this paper, many conserved target families were found to have conserved sequences flanking their miRNA binding-sites. Given these flanking sequences are enriched in homologues corresponding to HE targets, it suggests they may be facilitating strong/functional MTIs.

Many of these conserved sequences are predicted to form RNA secondary structures; *ARF10*, *ARF17*, *GAMYB*, *HAM1*, *IPS1*, *ARF2*, *ARF3* and *ARF4* (Figures 4-7; Zheng et al., 2017; Wong et al., 2018). The pattern of conservation of many of these regions to consistent with the formation of RNA secondary structures; the nucleotides of predicted RNA loops are highly variable, whilst the nucleotides predicted to form RNA stems are conserved, including instances of substitutions that maintained base pairing to form the predicted RNA stems.

Curiously, many of the conserved flanking sequences identified were predicted to base-pair with the miRNA binding-site. Although in the first instance a highly structured miRNA binding-site may seem counterintuitive, as accessibility may attenuate regulation, an *in vivo* assessment of RNA structure of miRNA binding-sites found them to be highly structured (Yang et al., 2020). Additionally, similar to the RNA stem-loops associated with the miR159 binding-site of *MYB33/65* (Zheng et al., 2017), the majority of conserved flanking sequences were located upstream and were predicted to base-pair with the 5‵ end of the miRNA binding-site, leaving the 3‵ end of the miRNA binding-site unbound (miR160:*ARF17*; miR171:*HAM1*; tasiARF:*ARF3*; tasiARF:*ARF4*). The 3‵-end region corresponds to the nucleotide positions most important for miRNA-mediated regulation (5‵end of the miRNA), as multiple studies have found that mismatches within this region preferentially attenuate regulation (Mallory et al., 2004; Schwab et al., 2005; Liu et al., 2014). Therefore, it is tempting to speculate that these structures are designed to promote accessibility to this region of the miRNA binding-site. For the miR160:*ARF10* and tasiARF:*ARF2* modules, conserved sequences were also found downstream of the binding site and were predicted to base-pair with the 3‵ region of the miRNA binding-site. Interestingly, in these cases, some of the 3‵ nucleotides of the binding site still coincided with the unpaired loop region of the predicted stem-loop which may be leading to an open conformation and greater accessibility. Alternatively, these predicted RNA structures may play a role in ribosome stalling as some studies have implicated RNA secondary structures in ribosome stalling in plants (Gawronski et al., 2018). Therefore, it may be that these structures cause the ribosome to stall and delay the completion of translation which therefore increases miRNA binding-site accessibility for increased silencing. Clearly, more work is needed here to determine whether these predicted RNA secondary structures exist *in vivo* and their function, if any, on miRNA-mediated regulation.

Finally, the conserved flanking sequences in the miRNA target, *ARF10*, was functionally demonstrated to be involved in miRNA-mediated regulation (Figure 9). Like in *MYB33*, the conserved flanking sequences also form a predicted RNA secondary structure. Furthermore, mutations to the flanking sequences, which was also predicted to alter the RNA secondary structure, attenuated miR160-mediated regulation. Thus, this further supports a role for RNA secondary structures flanking the miRNA binding in the miRNA-mediated regulation of some targets. Functional testing of these features in other miRNA targets may further shed light on the role factors beyond complementarity play in miRNA-mediated regulation.

## MATERIAL AND METHODS

### Bioinformatics workflow to identify HE and LE targets across species

Mature miRNA sequences for all species were retrieved from miRBase v22 (Kozomara et al., 2019). Where multiple isomiRs were found, the isomiR with the highest abundance found on a plant next-generation sequencing database (https://mpss.danforthcenter.org.) was used (Nakano et al., 2020). The isomiR sequences analysed can be found in Table S1. Targets were predicted using psRNATarget v2 (Dai et al., 2018). Default settings were used for analysis except the expectation score which was decreased to 3 for all miRNAs except miR167, miR398 and miR408. An expectation score of 5 was used for these miRNAs as their genes from the VAT set exceeds an expectation score of 3. The resulting predicted targets were then analysed using Whole-Degradome-based Plant MicroRNA-Target Interaction Analysis Server (WPMIAS) (Fei et al., 2020). The “Advanced II” > “Use psRNATarget predicted results directly” option was used for analysis by WPMIAS for all miRNAs. Default settings were used for all miRNAs except for miR162, miR396, miR398 and miR408 where “Offset from spliced position (nt)” was set to 1 as the previously validated targets of these miRNAs can only be identified at this setting.

The transcriptome libraries from psRNATarget and WPMIAS used for analysis can be found on Table S2. Degradome data retrieved from WPMIAS was then used as input and analysed using TRUEE to identify HE and LE targets. Analysis by TRUEE was performed at a *Cleavage Tag Abundance* of ≥ 5 TP10M, *Library % Cut-off* of 20% and a *Target Category* of both Category 1 and 2 targets. R script used for this analysis is accessible on the Open Science Framework page for this project https://osf.io/7anb9/. Target Categories as defined in WPMIAS were used in this study (Fei et al., 2020).

### Quantifying sequence conservation and RNA secondary structure prediction

For each target gene twenty sequences from diverse land plants were retrieved from nBLAST using the *A. thaliana* sequence as input. Species diversity was achieved by choosing species ranging across major taxonomic divisions where homologues were available. Taxonomic divisions were eudicots-rosids, eudicots-asterids, eudicots-ranunculids, monocots, *A. trichopoda*, gynmnosperms, lycophytes and bryophytes. One sequence was chosen for each species with the highest sequence identity of the whole gene. Only up to two mismatches in the binding site were allowed for each sequence. Default settings were used for BLASTn.

Sequences were aligned using Multiple Alignment using Fast Fourier Transform (MAFFT) with default settings (Katoh and Standley, 2013). Conservation was determined using phyloP (phylogenetic p-values) using LRT in “CON” conservation mode to measure slower than neutral evolution (Pollard et al., 2010) where a positive phyloP score denotes conservation. phyloP scores (p-values) were generated using Phylogenetic Analysis with Space/Time Models (rPHAST) (Hubisz et al., 2011). As input into rPHAST, phylogenetic trees were generated using Simple Phylogeny to fit phylogenetic tree to the alignment and determine a neutral model (https://www.ebi.ac.uk/Tools/phylogeny/simple_phylogeny/) (Larkin et al., 2007; Goujon et al., 2010; McWilliam et al., 2013, Madeira et al., 2019). The neutral model is changes of the sequence under neutral genetic drift. Hence by comparing the substitution rate at a particular nucleotide position to the neutral model, whether this nucleotide is conserved or undergoing accelerated substitution can be determined. phyloP scores were further adjusted for FDR using the “BH” method (Benjamin and Hochberg, 1995). In this study, Individual nucleotide positions were only considered conserved if they possessed an FDR-adjusted phyloP score of ≥ 1.0. Sequences were considered conserved if conserved nucleotides occurred ≥4 in a row. The R script used to calculate phyloP score is accessible on the Open Science Framework page for this project https://osf.io/7anb9/.

### Identification of the presence of conserved sequence in HE and LE targets across species

Having identified conserved sequences flanking the miRNA binding-site, the HE and LE targets of the primary target family were then analysed for the presence of these conserved sequences. Transcript sequences used to identify the presence of the conserved sequences for HE and LE targets of miRNA:target modules [miR159:*MYB33*; miR160:*ARF10* and *ARF17*; miR171:*HAM*; miR319:*TCP2* and*TCP4*; miR396:*GRF3*; tasiARF:*ARF2*, *ARF3* and *ARF4*] were retrieved from transcriptomes downloaded from the Genome portal of the Department of Energy Joint Genome Institute (Grigoriev et al., 2012; Nordberg et al., 2014) (https://phytozome.jgi.doe.gov/pz/portal.html) (Goodstein et al., 2012) (Table S3). The presence of the conserved sequences was identified using an in-house R script which is accessible on the Open Science Framework page for this project https://osf.io/7anb9/. The workflow is described in Figure S12).

### Data visualization

MSAs were visualised using Jalview (Waterhouse et al., 2009). The consensus sequence around the binding site, including the conserved sequences, was used to generate sequence logos using WebLogo (Crooks et al., 2004). The consensus RNA secondary structure was analysed using RNAalifold (Bernhart et al., 2008) from a sequence window consisting of ∼50 nt upstream and downstream of the binding site (50 nt + 21 nt + 50 nt = 121 nt window). This window was extended to approximately 100 nt upstream and downstream of the binding site for miR171:*HAM* where the conservation of sequences flanking the binding site appeared to extend beyond a window of 121 nt. Default parameters were used except temperature which was set at 22°C to generally reflect plant growth temperatures. T-plots of miRNA targets were adapted from WPMIAS (Fei et al., 2020). All graphs were generated using the R package, ggplot2, except for the pie charts which were generated using Excel.

### PANTHER ID acquisition

PANTHER IDs, which were used to sort HE and LE targets into gene families, from Phytozome v12 via Phytomine, the InterMine interface to Phytozome (https://phytozome.jgi.doe.gov/phytomine/begin.do) (Goodstein et al., 2012; Smith et al., 2012; Mi et al., 2013). Phytomine was accessed using IntermineR, an R package providing an R interface with InterMine-Powered Databases, and was incorporated into the R-script (Kyritsis et al., 2019). Genes with no associated PANTHER ID are excluded from analysis and is the cause of discrepancy in the numbers between analysis using (Figure 2) and not using (Figure 1) PANTHER IDs. Analysis was performed using Phytozome v12.

### Generation of Arabidopsis transformants expressing the *ARF10* variant transgenes

The *ARF10* gene sequence was amplified from genomic DNA using primers containing *attb1* and *attb2* sites (Table S4) using KOD Hot Start DNA Polymerase (Novagen™) and cloned into the donor vector, pDONOR/Zeo using the Gateway™ BP Clonase™ II enzyme mix (Invitrogen™) to produce an *ARF10* (henceforth *ARF10-WT*) entry clone. The *ARF10-FM* and *rmARF10* entry clones were generated by using primers (Table S4) to introduce mutations into the *ARF10-WT* entry clone using site-directed mutagenesis as previously described (Liu and Naismith, 2009), and were confirmed using restriction enzyme digestion analysis and DNA sequencing. The entry clones were then sub-cloned into the destination vector, pGWB602Ω (Nakamura et al., 2010), using the Gateway™ LR Clonase™ II enzyme mix (Invitrogen™) to generate the *ARF10-WT*, *ARF10-FM* and *rmARF10* expression clones, and confirmed clones were transformed into a GV3101 strain of *A. tumefaciens* by electroporation (Hellens et al., 2000) on LB agar plates containing 50 μg/mL Rifamycin, 25 μg/mL Gentamicin and 50 μg/mL Spectinomycin. *Arabidopsis thaliana* ecotype Columbia-0 (Col-0) plants were then transformed with each *ARF10* variant as previously described (Clough and Bent, 1998). Primary transformants were identified at 6-7 days old and transplanted onto soil.

### RNA secondary structure prediction

The consensus RNA secondary structure was analysed using RNAfold (Hofacker et al., 1994; Turner et al., 2009) from a sequence window consisting of approximately 50 nt upstream and downstream of the binding-site (100 nt + 21 nt + 100 nt = 221 nt window). Default parameters were used except temperature which was set at 22°C to reflect Arabidopsis growth temperatures.

### Statistical analysis

ANOVA was used to analyse the average Expectation score between HE targets and LE targets. Plant morphological phenotyping results were analysed using Pearson’s Chi-square test.

## AUTHOR CONTRIBUTIONS AND ACKNOWLEDGMENTS

G.Y.W. and A.A.M. designed the project. G.Y.W. performed the experiments. G.Y.W. and A.A.M. wrote the paper. We gratefully acknowledge the support of Zhi-Ping Feng from the ANU Bioinformatics Consultancy at the Australian National University. G.Y.W. was supported by an Australian Government Research Training Program RTP Scholarship.

## REFERENCES

Abdel-Ghany, S. E., and Pilon, M. (2008). MicroRNA-mediated systemic down-regulation of copper protein expression in response to low copper availability in Arabidopsis. Journal of Biological Chemistry, 283, 15932–15945.

Allen, E., Xie, Z., Gustafson, A. M., and Carrington, J. C. (2005). microRNA-directed phasing during trans-acting siRNA biogenesis in plants. Cell, 121, 207–221.

Axtell, M. J., and Meyers, B. C. (2018). Revisiting criteria for plant microRNA annotation in the era of big data. The Plant Cell, 30, 272–284.

Bari, A., Orazova, S., and Ivashchenko, A. (2013). miR156- and miR171-binding sites in the protein-coding sequences of several plant genes. BioMed Research International, 2013, 307145.

Benjamini, Y., and Hochberg, Y. (1995). Controlling the false discovery rate: a practical and powerful approach to multiple testing. Journal of the Royal Statistical Society. Series B (Methodological), 57, 289–300.

Bernhart, S. H., Hofacker, I. L., Will, S., Gruber, A. R., and Stadler, P. F. (2008). RNAalifold: improved consensus structure prediction for RNA alignments. BMC Bioinformatics, 9, 474.

Bonnet, E., He, Y., Billiau, K., and Van de Peer, Y. (2010). TAPIR, a web server for the prediction of plant microRNA targets, including target mimics. Bioinformatics, 26, 1566–1568.

Brousse, C., Liu, Q., Beauclair, L., Deremetz, A., Axtell, M. J., and Bouché, N. (2014). A non-canonical plant microRNA target site. Nucleic Acids Research, 42, 5270–5279.

Buscaglia, L. E., and Li, Y. (2011). Apoptosis and the target genes of microRNA-21. Chinese Journal of Cancer, 30, 371–380.

Chauhan, S., Yogindran, S., and Rajam, M. V. (2017). Role of miRNAs in biotic stress reactions in plants. Indian Journal of Plant Physiology, 22, 514–529.

Chen, Y., and Zhang, L. (2017). Members of the microRNA-200 family are promising therapeutic targets in cancer. Experimental and Therapeutic Medicine, 14, 10–17.

Chorostecki, U., Crosa, V. A., Lodeyro, A. F., Bologna, N. G., Martin, A. P., Carrillo, N., Schommer, C., and Palatnik, J. F. (2012). Identification of new microRNA-regulated genes by conserved targeting in plant species. Nucleic Acids Research, 40, 8893–8904.

Clough, S. J., and Bent, A. F. (1998). Floral dip: a simplified method for Agrobacterium - mediated transformation of *Arabidopsis thaliana*. The Plant Journal, 16, 735–743.

Crooks, G. E., Hon, G., Chandonia, J.-M., and Brenner, S. E. (2004). WebLogo: a sequence logo generator. Genome Research, 14, 1188–1190.

Dai, X., Zhuang, Z., and Zhao, P. X. (2018). PsRNATarget: A plant small RNA target analysis server (2017 release). Nucleic Acids Research, 46(W1), W49–W54.

Debernardi, J. M., Rodriguez, R. E., Mecchia, M. A., and Palatnik, J. F. (2012). Functional specialization of the plant miR396 regulatory network through distinct microRNA-target interactions. PLoS Genetics, 8, e1002419

Deveson, I., Li, J., and Millar, A. A. (2013). MicroRNAs with analogous target complementarities perform with highly variable efficacies in Arabidopsis. FEBS Letters, 587, 3703–3708.

Fahlgren, N., Montgomery, T. A., Howell, M. D., Allen, E., Dvorak, S. K., Alexander, A. L., and Carrington, J. C. (2006). Regulation of *AUXIN RESPONSE FACTOR3* by TAS3 ta-siRNA Affects Developmental Timing and Patterning in Arabidopsis. Current Biology 16, 939–944.

Fei, Y., Mao, Y., Shen, C., Wang, R., Zhang, H., and Huang, J. (2020). WPMIAS: Whole-degradome-based Plant MicroRNA–target Interaction Analysis Server. Bioinformatics, 36, 1937–1939.

Franco-Zorrilla, J. M., Valli, A., Todesco, M., Mateos, I., Puga, M. I., Rubio-Somoza, I., Leyva, A., Weigel, D., García, J. A., and Paz-Ares, J. (2007). Target mimicry provides a new mechanism for regulation of microRNA activity. Nature Genetics, 39, 1033–1037.

Gawroński, P., Jensen, P. E., Karpiński, S., Leister, D., and Scharff, L. B. (2018). Pausing of Chloroplast Ribosomes Is Induced by Multiple Features and Is Linked to the Assembly of Photosynthetic Complexes. Plant Physiology, 176, 2557–2569.

Goodstein, D. M., Shu, S., Howson, R., Neupane, R., Hayes, R. D., Fazo, J., Mitros, T., Dirks, W., Hellsten, U., Putnam, N., and Rokhsar, D. S. (2012). Phytozome: a comparative platform for green plant genomics. Nucleic Acids Research, 40(D1), D1178–D1186.

Goujon, M., McWilliam, H., Li, W., Valentin, F., Squizzato, S., Paern, J., and Lopez, R. (2010). A new bioinformatics analysis tools framework at EMBL-EBI. Nucleic Acids Research, 38, W695–W699.

Grigoriev, I. V., Nordberg, H., Shabalov, I., Aerts, A., Cantor, M., Goodstein, D., Kuo, A., Minovitsky, S., Nikitin, R., Ohm, R. A., Otillar, R., Poliakov, A., Ratnere, I., Riley, R., Smirnova, T., Rokhsar, D., and Dubchak, I. (2012). The Genome Portal of the Department of Energy Joint Genome Institute. Nucleic Acids Research, 40(D1), 26–32.

Gu, W., Wang, X., Zhai, C., Xie, X., and Zhou, T. (2012). Selection on synonymous sites for increased accessibility around miRNA binding sites in plants. Molecular Biology and Evolution, 29, 3037–3044.

Hellens, R. P., Allan, A. C., Friel, E. N., Bolitho, K., Grafton, K., Templeton, M. D., Karunairetnam, S., Gleave, A. P., and Laing, W. A. (2005). Transient expression vectors for functional genomics, quantification of promoter activity and RNA silencing in plants. Plant Methods, 1, 13.

Hofacker, I. L., Fontana, W., Stadler, P. F., Bonhoeffer, L. S., Tacker, M., and Schuster, P. (1994). Fast folding and comparison of RNA secondary structures. Monatshefte Für Chemie / Chemical Monthly, 125, 167–188.

Hubisz, M. J., Pollard, K. S., Siepel, A. (2011). PHAST and RPHAST: phylogenetic analysis with space/time models. Briefings in Bioinformatics, 12, 41–51.

Jones-Rhoades, M. W., and Bartel, D. P. (2004). Computational Identification of Plant MicroRNAs and Their Targets, Including a Stress-Induced miRNA. Molecular Cell, 14, 787–799.

Jones-Rhoades, M. W., Bartel, D. P., and Bartel, B. (2006). MicroRNAs and Their Regulatory Roles in Plants. Annual Review in Plant Biology, 57, 19–53.

Jones-Rhoades, M. W. (2012). Conservation and divergence in plant microRNAs. Plant Molecular Biology, 80, 3–16.

Katoh, K., and Standley, D. M. (2013). MAFFT Multiple Sequence Alignment Software Version 7: Improvements in Performance and Usability. Molecular Biology and Evolution, 30, 772–780.

Kawashima, C. G., Yoshimoto, N., Maruyama-Nakashita, A., Tsuchiya, Y. N., Saito, K., Takahashi, H., and Dalmay, T. (2009). Sulphur starvation induces the expression of microRNA-395 and one of its target genes but in different cell types. The Plant Journal, 57, 313–321.

Kozomara, A., Birgaoanu, M., and Griffiths-Jones, S. (2019). miRBase: from microRNA sequences to function. Nucleic Acids Research, 47(D1), D155–D162.

Kyritsis, K. A., Wang, B., Sullivan, J., Lyne, R., and Micklem, G. (2019). InterMineR: an R package for InterMine databases. Bioinformatics, 35, 3206–3207.

Larkin, M. A., Blackshields, G., Brown, N. P., Chenna, R., McGettigan, P. A., McWilliam, H., Valentin, F., Wallace, I. M., Wilm, A., Lopez, R., Thompson, J. D., Gibson, T. J., and Higgins, D. G. (2007). Clustal W and Clustal X version 2.0. Bioinformatics, 23, 2947– 2948.

Li, F., Zheng, Q., Vandivier, L. E., Willmann, M. R., Chen, Y., and Gregory, B. D. (2012). Regulatory impact of RNA secondary structure across the Arabidopsis transcriptome. The Plant Cell, 24, 4346–4359.

Li, J.-F., Chung, H. S., Niu, Y., Bush, J., McCormack, M., and Sheen, J. (2013). Comprehensive protein-based artificial microRNA screens for effective gene silencing in plants. The Plant Cell, 25, 1507–1522.

Li, J., Reichel, M., Li, Y., and Millar, A. A. (2014). The functional scope of plant microRNA-mediated silencing. Trends in Plant Science, 19, 750–756.

Liang, G., Yang, F., and Yu, D. (2010). MicroRNA395 mediates regulation of sulfate accumulation and allocation in Arabidopsis thaliana. The Plant Journal, 62, 1046–1057.

Lin, W.-Y., Lin, Y.-Y., Chiang, S.-F., Syu, C., Hsieh, L.-C., and Chiou, T.-J. (2018). Evolution of microRNA827 targeting in the plant kingdom. New Phytologist, 217, 1712–1725.

Liu, P. P., Montgomery, T. A., Fahlgren, N., Kasschau, K. D., Nonogaki, H., and Carrington, J. C. (2007). Repression of AUXIN RESPONSE FACTOR10 by microRNA160 is critical for seed germination and post-germination stages. The Plant Journal, 52, 133–146.

Liu, H., and Naismith, J. H. (2008). An efficient one-step site-directed deletion, insertion, single and multiple-site plasmid mutagenesis protocol. BMC Biotechnology, 8, 91.

Liu, Q., Wang, F., and Axtell, M. J. (2014). Analysis of complementarity requirements for plant MicroRNA targeting using a *Nicotiana benthamiana* quantitative transient assay. The Plant Cell, 26, 741–753.

Lustig, Y., Barhod, E., Ashwal-Fluss, R., Gordin, R., Shomron, N., Baruch-Umansky, K., Hemi, R., Karasik, A., and Kanety, H. (2014). RNA-Binding Protein PTB and MicroRNA-221 Coregulate AdipoR1 Translation and Adiponectin Signaling. Diabetes, 63, 433–445.

Madeira, F., Park, Y. mi, Lee, J., Buso, N., Gur, T., Madhusoodanan, N., Basutkar, P., Tivey, A. R. N., Potter, S. C., Finn, R. D., and Lopez, R. (2019). The EMBL-EBI search and sequence analysis tools APIs in 2019. Nucleic Acids Research, 47(W1), W636–W641.

Mallory, A. C., Reinhart, B. J., Jones-Rhoades, M. W., Tang, G., Zamore, P. D., Barton, M. K., and Bartel, D. P. (2004). MicroRNA control of PHABULOSA in leaf development: importance of pairing to the microRNA 5‵ region. The EMBO Journal, 23, 3356–3364.

Mallory, A. C., Bartel, D. P., and Bartel, B. (2005). MicroRNA-Directed Regulation of Arabidopsis AUXIN RESPONSE FACTOR17 Is Essential for Proper Development and Modulates Expression of Early Auxin Response Genes. Development, 17, 1–16.

McWilliam, H., Li, W., Uludag, M., Squizzato, S., Park, Y. M., Buso, N., Cowley, A. P., and Lopez, R. (2013). Analysis Tool Web Services from the EMBL-EBI. Nucleic Acids Research, 41, W597–W600.

Mi, H., Muruganujan, A., Casagrande, J. T., and Thomas, P. D. (2013). Large-scale gene function analysis with the PANTHER classification system. Nature Protocols, 8, 1551– 1566.

Millar, A. A., and Gubler, F. (2005). The Arabidopsis GAMYB-Like Genes, MYB33 and MYB65, Are MicroRNA-Regulated Genes That Redundantly Facilitate Anther Development. The Plant Cell, 17, 705–721.

Millar, A. A., Lohe, A., and Wong, G. (2019). Biology and Function of miR159 in Plants. Plants (Basel, Switzerland), 8, 255.

Nakamura, S., Mano, S., Tanaka, Y., Ohnishi, M., Nakamori, C., Araki, M., Niwa, T., Nishimura, M., Kaminaka, H., Nakagawa, T., Sato, Y., and Ishiguro, S. (2010). Gateway binary vectors with the bialaphos resistance gene, bar, as a selection marker for plant transformation. Bioscience, Biotechnology, and Biochemistry, 74, 1315–1319.

Nakano, M., McCormick, K., Demirci, C., Demirci, F., Gurazada, S. G. R., Ramachandruni, D., Dusia, A., Rothhaupt, J. A., and Meyers, B. C. (2020). Next-Generation Sequence Databases: RNA and Genomic Informatics Resources for Plants. Plant Physiology, 182, 136–146.

Naya, L., Paul, S., Valdés-López, O., Mendoza-Soto, A. B., Nova-Franco, B., Sosa-Valencia, G., Reyes, J. L., and Hernández, G. (2014). Regulation of Copper Homeostasis and Biotic Interactions by MicroRNA 398b in Common Bean. PLoS ONE, 9, e84416.

Nordberg, H., Cantor, M., Dusheyko, S., Hua, S., Poliakov, A., Shabalov, I., Smirnova, T., Grigoriev, I. V., and Dubchak, I. (2014). The genome portal of the Department of Energy Joint Genome Institute: 2014 updates. Nucleic Acids Research, 42, D26–D31.

Palatnik, J. F., Wollmann, H., Schommer, C., Schwab, R., Boisbouvier, J., Rodriguez, R., Warthmann, N., Allen, E., Dezulian, T., Huson, D., Carrington, J. C., and Weigel, D. (2007). Sequence and Expression Differences Underlie Functional Specialization of Arabidopsis MicroRNAs miR159 and miR319. Developmental Cell, 13, 115–125.

Pfeffer, S. R., Yang, C. H., and Pfeffer, L. M. (2015). The Role of miR-21 in Cancer. Drug Development Research, 76, 270–277.

Pilon, M. (2017). The copper microRNAs. New Phytologist, 213, 1030–1035.

Pollard, K. S., Hubisz, M. J., Rosenbloom, K. R., and Siepel, A. (2010). Detection of nonneutral substitution rates on mammalian phylogenies. Genome Research, 20, 110–121.

Reichel, M., and Millar, A. A. (2015). Specificity of plant microRNA target MIMICs: Cross-targeting of miR159 and miR319. Journal of Plant Physiology, 180, 45–48.

Rhoades, M. W., Reinhart, B. J., Lim, L. P., Burge, C. B., Bartel, B., and Bartel, D. P. (2002). Prediction of plant microRNA targets. Cell, 110, 513–520.

Schiefthaler, U., Balasubramanian, S., Sieber, P., Chevalier, D., Wisman, E., and Schneitz, K. (1999). Molecular analysis of NOZZLE, a gene involved in pattern formation and early sporogenesis during sex organ development in Arabidopsis thaliana. Proceedings of the National Academy of Sciences of the United States of America, 96, 11664–11669.

Schwab, R., Palatnik, J. F., Riester, M., Schommer, C., Schmid, M., and Weigel, D. (2005). Specific effects of microRNAs on the plant transcriptome. Developmental Cell, 8, 517– 527.

Smith, R. N., Aleksic, J., Butano, D., Carr, A., Contrino, S., Hu, F., Lyne, M., Lyne, R., Kalderimis, A., Rutherford, K., Stepan, R., Sullivan, J., Wakeling, M., Watkins, X., and Micklem, G. (2012). InterMine: a flexible data warehouse system for the integration and analysis of heterogeneous biological data. Bioinformatics, 28, 3163–3165.

Song J, Ouyang Y, Che J, Li X, Zhao Y, Yang K, Zhao X, Chen Y, Fan C and Yuan W (2017) Potential Value of miR-221/222 as Diagnostic, Prognostic, and Therapeutic Biomarkers for Diseases. Frontiers in Immunology, 8, 56.

Sun, Y.-H., Lu, S., Shi, R., and Chiang, V. L. (2011). Computational Prediction of Plant miRNA Targets. RNAi and Plant Gene Function Analysis: Methods and Protocols (H. Kodama and A. Komamine (Eds.); pp. 175–186). Humana Press.

Sunkar, R., Kapoor, A., and Zhu, J.-K. (2006). Posttranscriptional Induction of Two Cu/Zn Superoxide Dismutase Genes in Arabidopsis Is Mediated by Downregulation of miR398 and Important for Oxidative Stress Tolerance. The Plant Cell, 18, 2051–2065.

Sunkar, R., Li, Y.-F., and Jagadeeswaran, G. (2012). Functions of microRNAs in plant stress responses. Trends in Plant Science, 17, 196–203.

Tang, J., and Chu, C. (2017). MicroRNAs in crop improvement: fine-tuners for complex traits. Nature Plants, 3, 17077.

Todesco, M., Rubio-Somoza, I., Paz-Ares, J., and Weigel, D. (2010). A collection of target mimics for comprehensive analysis of microRNA function in *Arabidopsis thaliana*. PLoS Genetics, 6, 1–10.

Turner, D. H., and Mathews, D. H. (2010). NNDB: the nearest neighbor parameter database for predicting stability of nucleic acid secondary structure. Nucleic Acids Research, 38(Database issue), D280–D282.

Waterhouse, A. M., Procter, J. B., Martin, D. M. A., Clamp, M., and Barton, G. J. (2009). Jalview Version 2—a multiple sequence alignment editor and analysis workbench. Bioinformatics, 25, 1189–1191.

Wong, G., Alonso-Peral, M., Li, B., Li, J., and Millar, A. A. (2018). MicroRNA MIMIC binding sites: Minor flanking nucleotide alterations can strongly impact MIMIC silencing efficacy in Arabidopsis. Plant Direct, 2, e00088.

Wong, G. Y., and Millar, A. A. (2022). TRUEE; a bioinformatic pipeline to define the functional microRNA targetome of Arabidopsis. The Plant Journal, 110, 1476–1492.

Yamasaki, H., Abdel-Ghany, S. E., Cohu, C. M., Kobayashi, Y., Shikanai, T., and Pilon, M. (2007). Regulation of Copper Homeostasis by Micro-RNA in Arabidopsis. Journal of Biological Chemistry, 282, 16369–16378.

Yang, M., Woolfenden, H. C., Zhang, Y., Fang, X., Liu, Q., Vigh, M. L., Cheema, J., Yang, X., Norris, M., Yu, S., Carbonell, A., Brodersen, P., Wang, J., and Ding, Y. (2020). Intact RNA structurome reveals mRNA structure-mediated regulation of miRNA cleavage in vivo. Nucleic Acids Research, 48, 8767–8781.

Zheng, Z., Reichel, M., Deveson, I., Wong, G., Li, J., and Millar, A. A. (2017). Target RNA Secondary Structure Is a Major Determinant of miR159 Efficacy. Plant Physiology, 174, 1764–1778.

